# Tracking Lesion Growth in the Field: Imaging and Deep Learning Reveal Components of Quantitative Resistance

**DOI:** 10.1101/2025.05.20.655031

**Authors:** Jonas Anderegg, Lukas Roth, Radek Zenkl, Bruce A. McDonald

## Abstract

- Measuring individual components of pathogen reproduction is key to understanding mechanisms underlying rate-reducing quantitative resistance (QR). Simulation models predict that lesion expansion plays a key role in seasonal epidemics of foliar diseases, but measuring lesion growth with sufficient precision and scale to test these predictions under field conditions has remained impractical.
- We used deep learning-based image analysis to track 6889 individual lesions caused by *Zymoseptoria tritici* on 14 wheat cultivars across two field seasons, enabling 27,218 precise and objective measurements of lesion growth in the field.
- Lesion appearance traits reflecting specific interactions between particular host and pathogen genotypes were consistently associated with lesion growth, whereas overall effects of host genotype and environment were modest. Both host cultivar and cultivar-by-environment interaction effects on lesion growth were highly significant and moderately heritable (*h*^*2*^ ≥ 0.40). After excluding a single outlier cultivar, a strong and statistically significant association between lesion growth and overall QR was found.
- Lesion expansion appears to be an important component of QR to STB in most—but not all—wheat cultivars, underscoring its potential as a selection target. By facilitating the dissection of individual resistance components, our approach can support more targeted, knowledge-based breeding for durable QR.

## Introduction

Most crop disease epidemics are polycyclic, with multiple rounds of infection occurring during a single growing season. In epidemics driven mainly by locally produced inoculum, epidemic progression depends largely on the time required for the pathogen to complete its life cycle and the rate of pathogen reproduction associated with each cycle (Willocquet *et al*., 2017). Septoria tritici blotch (STB), caused by the fungus *Zymoseptoria tritici*, is one of the most damaging foliar diseases of wheat worldwide (Savary *et al*., 2019) and provides a prominent example of a disease characterized by locally driven polycyclic epidemics. At the field scale, STB epidemics are fueled by mass production of asexual conidia that are dispersed over short distances via rain splash (Suffert & Sache, 2011; Karisto *et al*., 2022). While the primary inoculum is composed mainly of genetically diverse airborne ascospores that immigrate from outside of a wheat field, the splash-dispersed conidia rapidly become the dominant driver of a field epidemic, which often begins as early as late autumn in winter crops (Suffert & Sache, 2011).

Resistance reactions of wheat cultivars exposed to field populations of *Z. tritici* are typically highly quantitative (e.g., Karisto *et al*., 2018). This quantitative resistance (QR) arises from a combination of factors, including the high genetic diversity found within field populations of *Z. tritici* (McDonald *et al*., 2022), the variable effectiveness across pathogen strains of individual major resistance genes encoding effector-triggered resistance (Meile *et al*., 2018, Meile *et al*., 2023; Langlands-Perry *et al*., 2023), and the deployment of diverse combinations of major resistance genes in different cultivars (Brown *et al*., 2015). Effector-triggered resistance (ETR) against *Z. tritici* primarily hinders stomatal penetration and early establishment of the fungus within host tissues (Battache *et al*., 2022, Battache *et al*., 2024; Alassimone *et al*., 2024), reducing infection frequency and, ultimately, the number of developing lesions. ETR is typically assessed using a single integrated disease severity measure such as percentage of symptomatic leaf area collected at a late stage of infection (e.g., Meile *et al*., 2018; Alassimone *et al*., 2024), based on experiments conducted in greenhouses using high inoculum concentrations and controlled environments that favor infection. This approach makes it difficult to disentangle the effects of individual components of disease development such as infection frequency, incubation period, latent period, or lesion growth rate, on overall disease progression (Karisto *et al*., 2019; Anderegg *et al*., 2024b). Yet, QR results not only from a reduced infection frequency but also from other factors influencing the rate of disease progression within each infection cycle. For example, for the septoria nodorum blotch (SNB) disease on wheat that shares many epidemiological processes with STB, Jeger *et al*. (1983) and Lancashire & Jones (1985) observed differences in various components of QR among wheat genotypes, including duration of incubation period and latent period, initial lesion size and growth rate, and spore production. In the same pathosystem, Adhikari *et al*. (2023) found major differences in field-measured lesion growth rates across different wheat cultivars, and lesion growth rates were closely related to the overall level of QR in those cultivars. Similar studies investigating these components of QR remain scarce for STB, with most works focusing on spore production and duration of latent and incubation periods under controlled conditions (e.g., Brokenshire, 1976; Simon & Cordo, 1997, 1998). Compared to ETR that primarily reduces infection frequency, the molecular mechanisms underlying variation in other components of QR remain poorly understood (Poland *et al*., 2009; Cowger & Brown, 2019).

*Z. tritici* has a long latent period, typically lasting 3-4 weeks under field conditions (Lovell *et al*., 2004; Armour *et al*., 2004) as well as specific environmental requirements for spore dispersal, germination, and infection. As a result, the pathogen typically undergoes only around 4-6 infection cycles during the spring and summer periods of most rapid crop growth in temperate Europe (Karisto et al., 2018). Other wheat diseases have shorter latent periods under field conditions [e.g., 6 days for SNB (Ficke *et al*., 2018), 5 days for leaf rust (Wójtowicz *et al*., 2017), and 5 days for tan spot (Jørgensen & Olsen, 2007)] that enable many more infection cycles in the same period of time. Therefore, the rate of disease progression and pathogen multiplication per cycle plays a critical role in shaping STB epidemics. Compared to spore dispersal in the canopy, disease progression on infected leaves within a single infection cycle is easier to quantify. Key components can be measured or estimated using relatively straightforward methods, including manual counts of lesions and fruiting bodies, lesion size measurements, or visual assessments of diseased leaf area over time (Shaner, 1973; Parlevliet, 1975, 1979; Magboul *et al*., 1992; Bernard *et al*., 2013; Adhikari *et al*., 2023). Among these components, lesion growth plays an important role in shaping STB disease progression over time. It proportionately increases inoculum production, and necrotic tissue becomes infectious almost immediately. Moreover, lesion growth contributes to disease intensification even when conditions are unfavorable for new infections or when susceptible leaf tissue is no longer available for new infections (Berger *et al*., 1997). Lesion growth also has important agronomic implications, as it directly reduces assimilate availability for crop growth and development.

Identifying components of QR that slow disease progression and reduce pathogen multiplication within each infection cycle could significantly enhance resistance breeding by enabling recombination of genetically and functionally distinct resistance mechanisms (Jeger *et al*., 1983; Willocquet *et al*., 2017; Anderegg *et al*., 2024b). Unfortunately, this concept is difficult to implement in practice because components of rate-reducing QR, besides being time-consuming to assess, exhibit poorly characterized host-pathogen-environment interactions, further complicating their unbiased measurement. For example, the duration of the latent period in STB is affected not only by pathogen and host genotype, but also by leaf temperature and leaf temperature variation (Bernard *et al*., 2013, Bernard *et al*., 2022). Similarly, SNB lesion growth rates under field conditions depend not only on host genotype, but also on temperature and genotype-by-temperature interactions (Adhikari *et al*., 2023). A quantitative understanding of these interactions is needed for the development of more accurate epidemiological models. However, more comprehensive data sets are needed to quantify the most important interactions that contribute to the frequently observed genotype-by-environment interactions in overall QR (e.g., Schilly *et al*., 2011).

Several studies have sought to replace tedious and subjective manual assessments of QR components with more objective and higher-throughput image-based approaches (Pelletier & Fry, 1989; Díaz-Lago *et al*., 2003; Behmann *et al*., 2018; Karisto *et al*., 2019; Anderegg *et al*., 2024b). Recent advances in imaging technology and image analysis, particularly end-to-end trainable deep neural networks, have enabled faster and more objective quantification of multiple disease-related traits at both the canopy and individual leaf levels directly in the field and under naturally fluctuating environmental conditions. This has allowed a significant increase in measurement throughput and enabled transformation of image-based methods into new phenomics tools capable of capturing detailed information on multiple disease-related traits based on a single measurement (Anderegg *et al*., 2023, 2024b; Zenkl *et al*., 2025). These new approaches allow precise and detailed characterization of developing disease symptoms within their real-world spatial context. Lesion area can be quantified at pixel-level accuracy for all symptoms present simultaneously, while shape parameters, fruiting body counts, densities, and spatial distributions, as well as lesion edge characteristics can also be analyzed. Contextual information, such as the total number of symptoms present on a leaf, or the occurrence of other foliar diseases can be extracted from the same measurement (Anderegg *et al*., 2024b).

The overall objective of this study was to utilize these novel image-based phenomics approaches to investigate drivers of lesion growth in wheat leaves infected with *Z. tritici* under natural field conditions. Specifically, we aimed to quantify the effect of the host, the environment, as well as properties of individual symptoms representing specific host-pathogen interactions on lesion growth rates. Our results shed new light on the predictability of lesion growth, the extent to which genetic and environmental factors determine lesion growth rates, and the relationship between lesion growth and overall QR in the wheat/*Z. tritici* pathosystem.

## Materials and Methods

### Field Experiments

An overview of data acquisition, processing, and analysis is given in Figure 1. Leaf-level development of STB was monitored in field experiments carried out in the wheat growing seasons of 2022-2023 and 2023-2024 at the ETH Research Station for Plant Sciences at Lindau-Eschikon, Switzerland (47.449 N, 8.682E, 520 m a.s.l.; soil type: eutric cambisol). The experiments were carried out on adjacent lots of the same site. Twenty registered European wheat cultivars with contrasting morphology and varying degrees of susceptibility to STB were used for the experiment. Although the level of QR was not the primary criterion for selecting cultivars, the chosen set represented the full range of resistance observed among 335 recent European wheat varieties. This is supported by a single-year visual assessment of QR (Karisto *et al*., 2018), which, though limited, offers a reasonable approximation of resistance variation.

**Figure 1.**
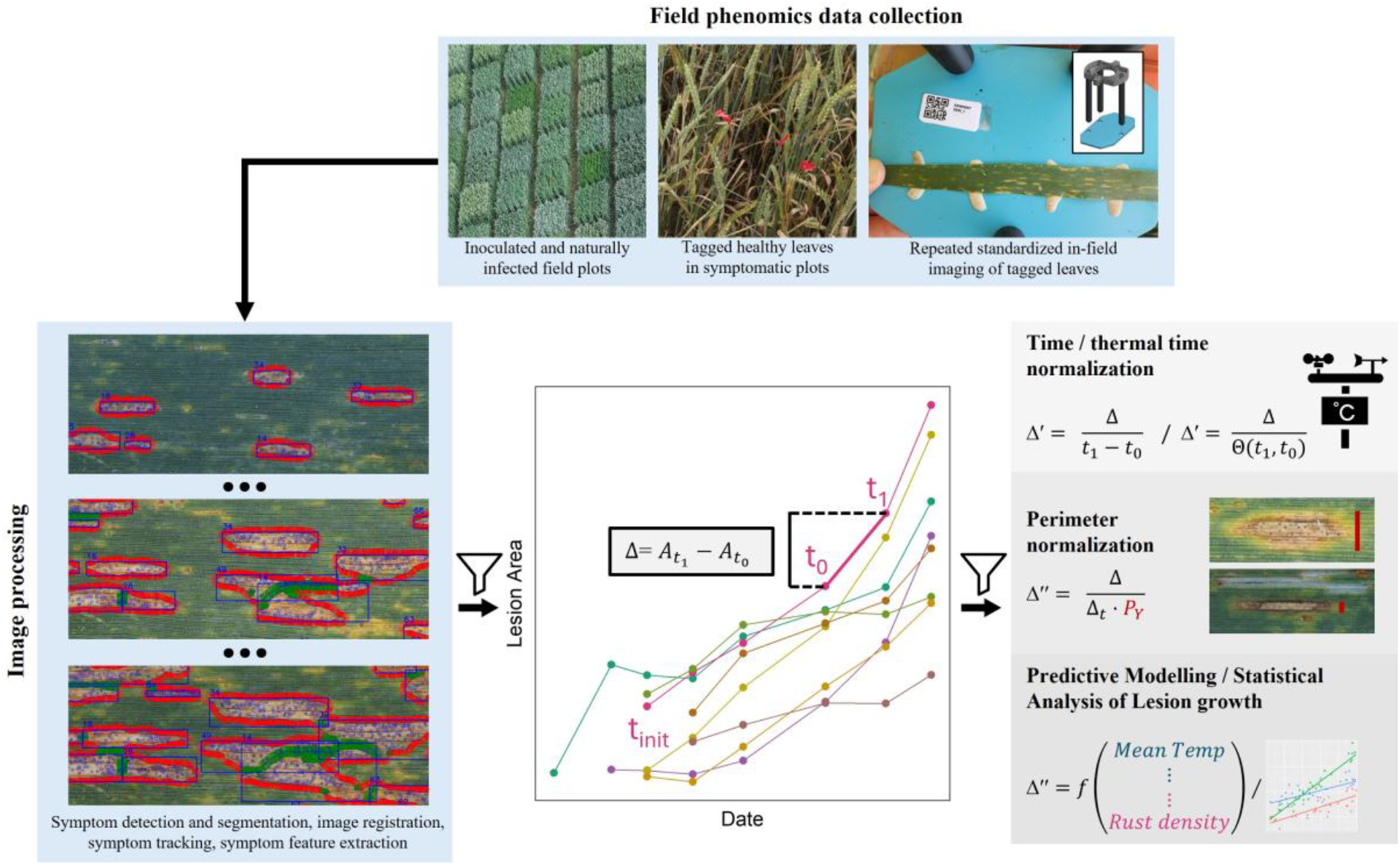
Flow diagram illustrating the steps involved in data acquisition, processing, and analysis. Image-based lesion growth measurements were combined with lesion- and leaf-level phenomics as well as meteorological data to investigate the dependency of lesion growth on host cultivar, environment, and general leaf-level context. For more details on data acquisition and image processing, refer to Anderegg *et al*. (2024) and Zenkl *et al*. (2025).

An overview of the cultivars is provided in Supplementary Table S1. Each cultivar was grown in 9 replicate plots sized 1 m × 1.7 m, of which three were assigned to each treatment. The 2-factorial experimental design (cultivars and treatments) was generated using the functions *findblks()* and *facDiGGer()* of the R-package ‘DiGGer’ (Coombes, 2009). The treatments were (i) an early fungicide application at jointing followed by a late artificial inoculation with a *Z. tritici* spore suspension at full flag leaf emergence (F_I2), (ii) no fungicide application but an early artificial inoculation at booting, when penultimate leaves were extending (0F_I1), and (iii) an early fungicide application at jointing without artificial inoculation (F_0I). In addition, one replicate plot per cultivar (three for the check cultivar ‘CH Claro’) was available where neither fungicide nor inoculum was applied (0F_0I). Artificial inoculations were applied to maximize the chances of being able to observe significant STB development in case environmental conditions would be unfavorable for the development of a natural epidemic. The fungicide used was “Input” (Bayer; a mixture of spiroxamine at 300 g/l and prothioconazole at 150 g/l), applied at 1.25 l/ha. Spore suspensions contained a mixture of 10 *Z. tritici* strains. For more details regarding the experimental design, refer to Anderegg *et al*. (2023). Air temperature and relative humidity were measured at 0.1 m above ground within the canopy of experimental wheat plots, using Campbell CS215 sensors (Campbell Scientific Inc., USA), installed in another field experiment conducted at the same site. A value was recorded every 10 min. In 2023, data from one plot was available, whereas in 2024, data from three sensors installed in different plots were averaged.

### Image sequences for monitoring STB development

Starting from three weeks after each inoculation, the uppermost leaves targeted by the inoculation (i.e., penultimate leaves or flag leaves) were monitored daily for STB. As soon as the first symptoms appeared, plots showing the highest incidence of fresh STB symptoms on the leaf layer of interest were selected for STB monitoring. Plot selection was based exclusively on STB incidence without reference to host genotype or treatment because measurements on non-symptomatic leaves would have been a waste of resources. In each selected plot, ten intact, fully developed, but still symptom-free leaves were tagged. The resulting sample distribution by genotype, treatment, and year is summarized in Supplementary Figure S1. In total, 386 and 646 single leaves were tagged and repeatedly imaged over multiple days at near-daily intervals in 2023 and 2024, respectively, following a previously described protocol (Anderegg *et al*., 2024b). Aligning with the two staggered inoculations that targeted penultimate leaves and flag leaves, leaf monitoring was conducted in two consecutive temporal batches (batch1– penultimate leaves, and batch2–flag leaves). From the resulting image data set, only images with STB as the dominant cause of necrosis were retained for further analysis. For each image series, a start and end date were defined, identifying the portion of the image series meeting this criterion, based on visual inspection. This led to the exclusion of many leaves with no or very few developing STB lesions, with severe yellow rust or brown rust infections, or with advanced physiological senescence. 392 leaves out of the total of 1032 were retained for further analysis. The resulting sample distribution by genotype, treatment, and year is summarized in Supplementary Figure S2. Most sampled leaves originated from plots that had received no fungicide application in combination with the first inoculation (0F_I1; Supplementary Figure S2A). However, 80 leaves (∼20%) originated from plots without artificial inoculation, thus representing natural infections.

### STB lesion growth measurement

Disease symptoms present on imaged leaves were detected and segmented using deep-learning-based key point detection and semantic segmentation models, respectively. Pre-existing models were fine-tuned for the present use case, leveraging additional annotated training data consisting of 258 annotated image patches sized 1024 × 1024 pixels (n = 211, newly annotated) or 1224 × 1224 pixels (n = 47, associated with Anderegg *et al*., 2024b). Symptom tracking in the series of aligned images based on area overlap enabled accurate quantification of the growth of each lesion over time. Details on image acquisition and alignment, disease detection and segmentation, and symptom tracking and characterization can be found in our previous work (Anderegg *et al*., 2022, Anderegg *et al*., 2024b; Zenkl *et al*., 2025).

Filtering the image data set at the leaf level did not eliminate all false positive detections of STB lesions. Therefore, additional filtering criteria were applied at the lesion level to exclude leaf damage resulting from other diseases or of unknown origin. The presence of pycnidia was used as a positive diagnostic marker for STB. Specifically, lesions that did not contain at least 3 pycnidia on average across measurements falling within 120 hours of their initial detection were excluded, as were lesions with a low pycnidia density (<0.0005 px^-1^). Lesions younger than 96 hours with a high average rust pustule density (>0.0003 px^-1^) across these early time points were also excluded, due to the risk that lesion borders or the entire lesion were influenced more by rust than by STB. Similarly, lesions with average pycnidia density below 1/3 of rust pustule density were removed, without reference to lesion age. Finally, all lesions that had obstructions to growth along more than 50% of their perimeter (i.e., neighboring lesions, leaf edges, insect damage, or edges of the region-of-interest) were excluded because they did not provide a solid basis for observation of lesion growth.

In agreement with previous observations under controlled environment conditions (Karisto *et al*., 2019), new lesions continued to emerge throughout the measurement period on most analyzed leaves, despite two separate artificial inoculation events being the most likely cause of synchronized infections on most leaves. Many lesions were still expanding when measurements were stopped mostly due to advanced leaf senescence. Accordingly, measurements did not capture the entire growth process for each lesion but rather captured some subset of the growth process that was more complete for early-emerging lesions and increasingly incomplete for later-emerging lesions with fewer repeat observations. Direct modelling of each measurement series separately as is typically done for more complete measurements of more predictable trait trajectories (e.g., plant height or senescence development of wheat cultivars; Roth *et al*. (2024); Anderegg *et al*., (2024a)) was therefore not feasible in the present scenario. Following Adhikari *et al*. (2023), we therefore converted *N* timepoint observations of lesion size to *N*-1 interval observations of lesion area growth between a given timepoint and its lag timepoint (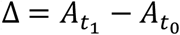; cf. Figure 1). To normalize for time, growth was divided by the duration of the interval (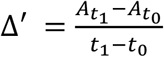; cf. Figure 1). Values outside the 1^st^ and 98^th^ percentiles of the resulting growth measurements distribution were removed to reduce the influence of extreme observations and enhance the robustness of subsequent analyses (see supplementary Figure S3). For ease of notation, the first timepoint of the interval (i.e., the lag timepoint) will be referred to as t_0_, whereas the second timepoint of the interval will be referred to as t_1_ throughout (Figure 1).

After all filtering, a total of 27,218 lesion area growth measurements were available, with two consecutive measurements taken on average (mean ± standard deviation) 28.2 ± 14.5 hours apart. These measurements came from 6,889 unique lesions, measured 3.95 ± 2.8 times over a period of 7.5 ± 3.7 days. As expected, lesion growth measurements varied by cultivar due to differences in QR, ranging from 139 measurements from 42 individual lesions on the resistant cultivar ‘ARINA’ to 4,927 measurements from 1,206 individual lesions on the susceptible cultivar ‘BORNEO’. An overview of unprocessed lesion size data is provided in Supplementary Figure S4.

The absolute increase in lesion area (in mm^2^) during a measurement interval is typically assumed to depend on a measure of lesion dimension at t_0_ such as lesion length, radius, or a direct measure of the lesion area, as reviewed by Berger *et al*. (1997). Depending on the chosen assumption, lesion growth over time is expected to follow either linear or exponential growth patterns. For example, Adhikari *et al*. (2023) assumed uniform growth along both axes of SNB lesions and therefore applied a square-root transformation to area measurements to obtain linearized growth rates in mm day^-1^. Here, we argue that lesion growth is not directly driven by lesion dimensions, but rather by fungal hyphal growth at the lesion perimeter. Accordingly, lesion growth should be some function of the length of that perimeter. The distinction between simplified measures of lesion dimension and a precise measurement of the perimeter length could be particularly relevant in cases where lesions are not of convex shape. Since growth of STB lesions occurs predominantly parallel to the major axis of the leaf along leaf veins (roughly corresponding to the x-axis of the image), the perimeter perpendicular to the major leaf axis should arguably be considered specifically (cf. Supplementary methods). Thus, to normalize for the purely geometric effect of lesion size on lesion area growth per unit of time, we expressed lesion area growth per unit of time relative to the length of the perimeter perpendicular to the major leaf axis that can support growth (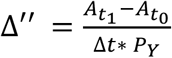, with *P*_*Y*_ denoting the length of the perimeter perpendicular to the major leaf axis; cf. Figure 1). This should also eliminate bias due to obstructions to growth along the lesion perimeter. All required information to derive this normalized measure of lesion growth can be derived from our image processing method (Anderegg *et al*., 2024b).

To examine the relationship between various factors other than environmental conditions (such as lesion size, lesion age, or measurement batch) and lesion growth, the duration of measurement intervals was converted from chronological time to effective thermal time based on measured temperature courses during each measurement interval. To normalize for thermal time, growth was then divided by the accumulated thermal time between t_0_ and t_1_ (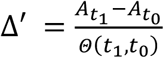, with *Θ*(*t*_1_, *t*_0_) denoting the accumulated thermal time between t_0_ and t_1_). A four-parameter beta function with the related cardinal temperatures and scaling factor estimated by Chaloner *et al*. (2019) for in-vitro hyphal growth were used for the conversion (Supplementary Figures S5 and S6). Due to the skewed (theoretically zero-bound) distribution of lesion growth data and heteroscedasticity (e.g., larger variance in lesion growth for larger compared to smaller lesions), quantile regression was preferred over mean-based modelling to examine pair-wise relationships between lesion growth and various factors such as lesion age or lesion area. Quantile regression was implemented using the functions *rq(), lprq()*, and *nlrq()* of the R-package ‘quantreg’ (Koenker *et al*., 2025). We assessed model fit using a pseudo-*R*^*2*^ metric for quantile regression, as proposed by Koenker & Machado (1999). This measure compares the sum of absolute residuals from the fitted model to that of an intercept-only model, providing an interpretable goodness-of-fit statistic.

### Predictive modelling of lesion growth and identification of most predictive features

Compared to traditional methods of assessing lesion growth, our newly developed phenomics tools offer a more comprehensive characterization of lesions and their context. We aimed to quantify the predictability of lesion growth based on this additional information for three main purposes: (i) to identify potential nuisance factors that need to be taken into account when modelling effects of host genotype and environment on lesion growth; (ii) to contextualize estimated effect sizes for key factors of interest (such as host genotype and environmental variables such as temperature) in terms of their contribution to the overall predictable variation; and (iii) to enhance functional understanding of the mechanics driving quantitative lesion growth under natural field conditions.

For each tracked lesion at each timepoint, a vector of features describing visual appearance of the lesion, the leaf it was located on, the measurement interval during which expansion occurred, and experimental design factors, was assembled. A description of all extracted features is provided in Supplementary Table S2. Details regarding feature extraction are reported in Supplementary methods and resulting feature distributions are shown in Supplementary Figure S7. The predictability of lesion growth during a measurement interval was then assessed by modelling lesion growth as a function of these features measured at t_0_, or during the interval itself (for interval features). As a baseline predictor, the mean of the response variable was used. Random forest (RF) regression models as implemented in the R-package ‘*ranger’* (Wright & Ziegler, 2017) were employed as predictive models, and tuned/trained via the ‘*caret’* package (Kuhn *et al*., 2021). To identify a subset of most informative yet relatively independent (un-correlated) features, we applied RF-based recursive feature elimination (RF-RFE) in multiple iterations, following a procedure described earlier (Anderegg *et al*., 2020). RFE is a feature selection method that iteratively removes the least important features, based on a nested cross-validation approach consisting of an inner loop for feature selection and an outer loop for model evaluation to prevent overfitting and ensure robustness (Guyon *et al*., 2002). Feature importance in each iteration was estimated via permutation. The importance of factor variables was calculated by summing up the importances of all factor levels obtained during one-hot-encoding of the corresponding factor variable. In the first iteration, RF-RFE was performed starting from the full model that involved all features. Features were assigned a rank corresponding to the RFE iteration during which they were eliminated. Based on these ranks, features that were highly redundant (correlated with another feature at r ≥ 0.925) were removed, dropping the feature with the higher average feature rank (indicating lower relevance for prediction). RF-RFE was then repeated starting from this reduced model. Redundant features (correlated with another feature at r ≥ 0.85) were again eliminated, dropping the less relevant feature. Finally, RF-RFE was performed starting from this further reduced feature set to finalize the selection of the most predictive and relatively uncorrelated features. To achieve acceptable runtime, only the ‘mtry’ hyper-parameter was optimized for each iteration and each subset, using 7-fold cross-validation, whereas ‘num.trees’ was set to 150, ‘min.node.size’ was set to 5, and all other parameters were set to their default values. Root mean squared error (RMSE) was used as the performance metric. It is important to note that high feature ranks (indicating a low relevance for growth prediction) as determined during this procedure do not necessarily indicate irrelevant features, as high ranks can also result from the corresponding factor being at least partially explained by one or a combination of other features in the data set. In contrast, consistently low feature ranks (indicating a high relevance for growth prediction) can indicate important features also in the presence of extensive multicollinearity in the data set. However, due to the inherent structure and imbalance in the dataset, as well as the partial confounding between variables (e.g., environmental conditions and measurement batches), insights from predictive modeling must be interpreted with caution and cannot replace a more thorough statistical analysis of the key predictors identified (see next section).

### Statistical analysis of cultivar and environmental effects on lesion growth

Effects of host cultivar, environmental variables, and cultivar-by-environment interactions on lesion growth were analyzed using a random regression approach. The models were specifically designed to estimate the main effects of host cultivar, mean interval temperature, and interval relative humidity, as well as the effects of interactions between host cultivar and both environmental variables, on lesion growth. They additionally incorporated the structure and hierarchical relationships within the data as well as key factors identified through feature selection (see above). These models thus provide a more comprehensive understanding of the factors contributing to lesion growth, beyond predictive ability. The base model was:

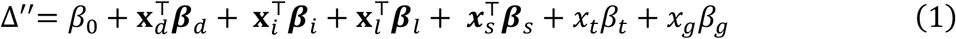

where Δ″ is the growth of a lesion in a specific interval, measured as the change in lesion area between consecutive time points per unit of time per unit of lesion perimeter (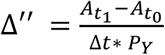; cf. Figure 1), *β*_0_ is the intercept, 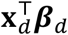 represents fixed effects of design-level covariates *x*_*d*_ (*d* = [year, batch]), 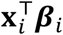 represents fixed effects of interval-level covariates *x*_*i*_ (*i* = [mean interval relative humidity, mean interval temperature]), 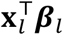 represents fixed effects of lesion-level covariates *x*_*l*_ (*l* = [lesion area, lesion age, maximum pycnidia distance from the lesion perimeter at t_0_]), 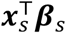 represents random effects of nested measurement levels *x*_*s*_ (*s* = [plot, leaf, lesion]) with *β*_*s*_ 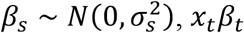 a random slope for age within lesions with *β*_*t*_ *∼ N*(0, *σ*^2^), and *x*_*g*_ *β*_*g*_ a random effect of genotype with *β*_*g*_ *∼ N*(0, *σ*^2^). Then, random regression effects for genotype-by-age, genotype-by-humidity, and genotype-by-temperature interactions on lesion growth were successively added to the baseline model. Thus, the final model was:

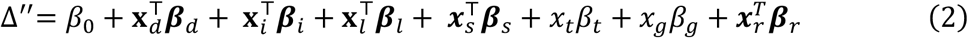

where 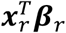 represents genotype-specific random regressions of lesion growth on lesion age, mean temperature, and mean relative humidity, modelled with a correlated variance-covariance structure ***β***_*r*_ *∼ N*(0, ***G*** ⊗ ***I***) where ***G*** is the 3×3 heterogeneous correlation matrix representing correlations between age, mean temperature, and mean relative humidity effects, ***I*** is the genotype identity matrix, and ⊗ the Kronecker product. This allowed for a joint estimation of the sensitivity of lesion growth on different host cultivars to multiple environmental gradients. Nested models were compared using the Bayesian information criterion and likelihood ratio tests. The models were fitted in R using ASReml-R (Butler *et al*., 2018).

To estimate the heritability of the host cultivar’s main effect on lesion growth as well as of the genotype-specific responses of lesion growth to temperature and relative humidity, we fitted model (2) for each individual experimental plot without including genotype information. Here, *x*_*g*_*β*_*g*_ thus represents the random effect of the plot and 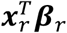 the plot-specific random regressions of lesion growth on lesion age, mean temperature, and mean relative humidity. Since models with a correlated variance-covariance structure did not converge, the responses were considered uncorrelated in this case, i.e., 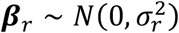. Then, the estimated plot-level regression slopes and the plot main effects on lesion growth were modelled as

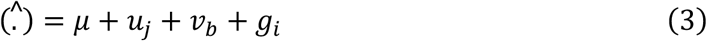

where 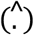 is either one of 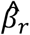 or 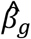, μ is a fixed intercept, *u*_*j*_ represents the fixed effects of the *j*^th^ year, *v*_*b*_ represents the fixed effects of the *b*^th^ batch, and *g*_*i*_ represents the random effect of the host genotypes *i*. Heritability was then estimated according to Cullis *et al*. (2006).

### Relationship between lesion growth and overall QR

The level of QR to STB of the cultivars included in the experiment was not known at the time of experimental design. Some genotypes were selected based on prior knowledge of their level of QR from (Karisto *et al*., 2018), but that information was partial, with a focus on late-season conditional disease intensity on infected leaves observed in one year. To obtain objective and precise estimates of QR for each genotype, we combined visual scoring of disease incidence with image-based assessments of conditional disease severity following a protocol described earlier (Anderegg *et al*., 2019). Assessments at five different time points across two years were aggregated. Data and statistical procedures are described in detail in Supplementary Methods. Relationships between lesion growth and components of overall QR were examined using Pearson correlation and linear regression.

## Results

### Weather patterns

Temperature, relative humidity, and rainfall patterns differed between the two experimental periods in 2023 and 2024 (Figure 2A). Including data from both years notably expanded the range of relative humidity values at which measurements were taken (Figure 2C), while the temperature range remained relatively constant (Figure 2B). A wider temperature range was achieved by measuring two consecutive batches of leaves, particularly in 2023 (Figure 2B). Variation in weather conditions was therefore significantly confounded with both year and batch, with relatively narrow temperature and relative humidity ranges observed within individual batches.

**Figure 2.**
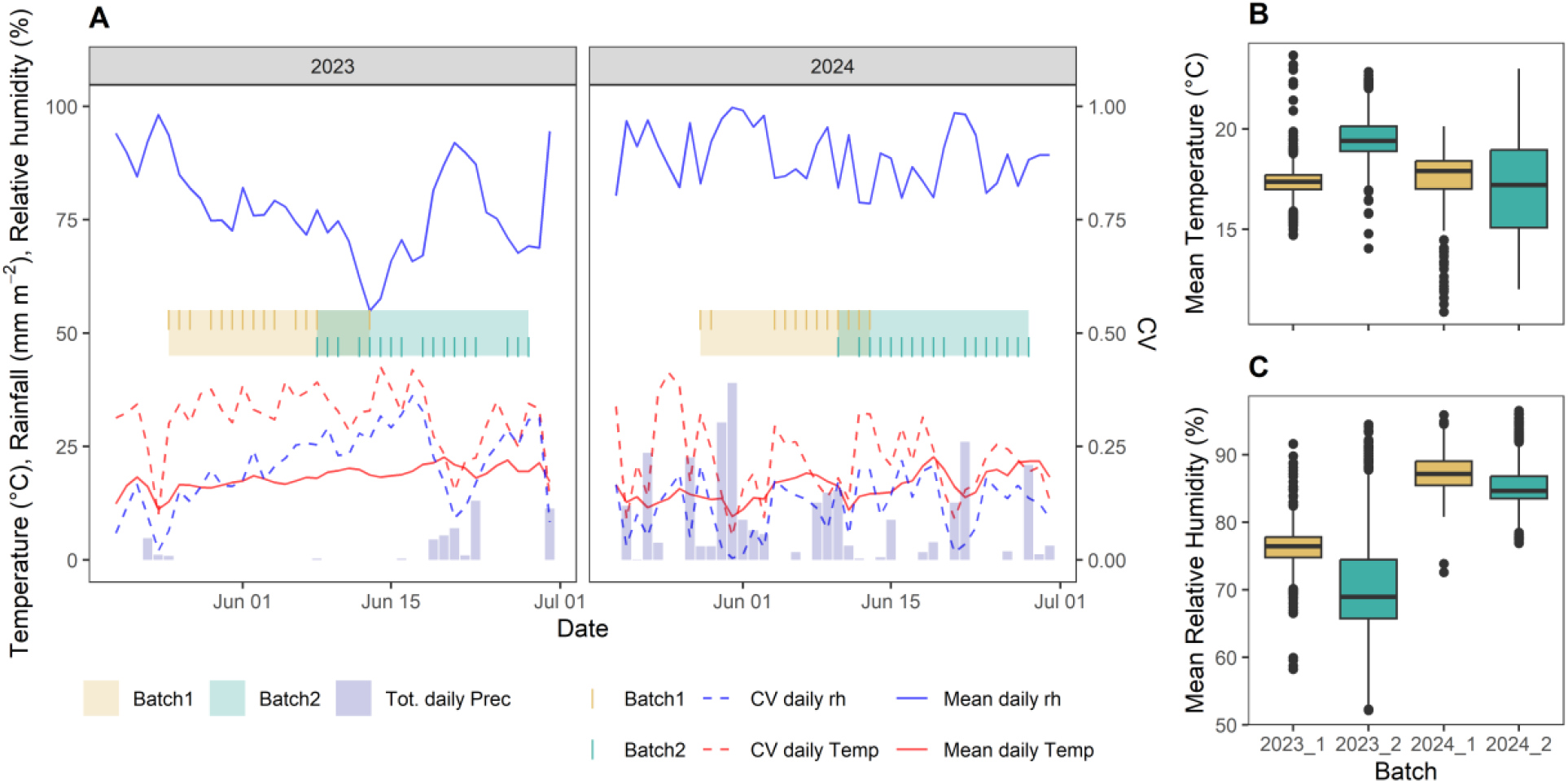
Environmental conditions during lesion growth measurement periods. (**A**) Weather conditions during the experimental periods of 2023 and 2024. Batches refer to experimental periods and to the leaf layer investigated (Batch1 – penultimate leaves, first cohort of lesions; Batch2 – flag leaves, second cohort of lesions). Small colored lines indicate individual measurement events. (**B**) Boxplot of mean temperatures recorded during measurement intervals. (**C**) Boxplots of mean relative humidity recorded during measurement intervals.

In general, temperatures were higher, and relative humidity was lower, during the 2023 measurement period (Figure 2A). Daily variability in temperature and relative humidity was higher in 2023, whereas there was much more rainfall during the measurement period in 2024. This resulted in a longer measurement gap at the end of May of that year (Figure 2A).

Despite the highlighted differences between years, weather conditions in both growing seasons were generally conducive to the development of STB. In 2024, disease incidence on flag leaves reached nearly 100% by mid-grain filling on many European cultivars tested at the experimental site, despite intense control measures (i.e., three fungicide treatments and a crop rotation; unpublished data). This is not uncommon at the study site (Karisto *et al*., 2018). In 2023, overall disease pressure appeared to be somewhat lower, but substantial natural infection was also observed and contributed to the data set analyzed herein (Supplementary Figure S2).

### Retrieving lesion growth from repeated lesion area measurements

Lesion area increased with lesion age, but the relationship varied considerably across individual lesions, host cultivars, and measurement batches (Figure 3). When aggregating all measurements, a nearly linear trend was observed between lesion age and lesion area (*R*^*2*^ = 0.35, *P* < 2.2 × 10^−16^, Supplementary Figure S8). Differences in lesion perimeter lengths at t_0_ accounted for up to 18% of the total variation in absolute lesion growth (expressed as mm^2^ per unit of thermal time) during the subsequent time interval (Supplementary Figure S9). The relationship was significantly non-linear for all perimeter dimension(s) considered, with all second-order polynomial terms highly significant (*P* < 2.2 × 10^−16^). As expected, lesion growth was more strongly related to the length of the perimeter in the direction perpendicular to the major leaf axis. Surprisingly, correcting the lesion perimeter lengths for obstructions to growth did not strengthen the correlations. Thus, all subsequent analysis used change in lesion area as normalized by the length of the perimeter perpendicular to the major leaf axis at t_0_ as the measure of lesion growth, unless specified otherwise.

**Figure 3.**
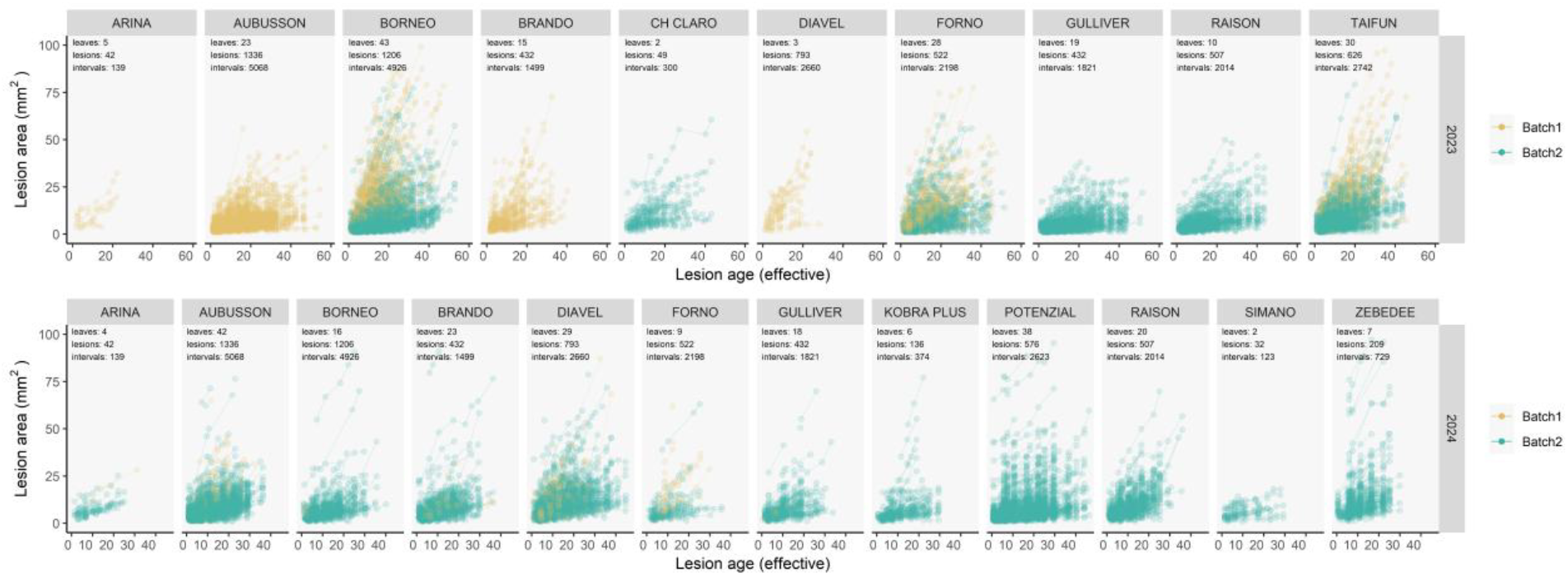
Lesion area as a function of effective lesion age in thermal time, separately for each host cultivar in each year. Lines connect repeat measurements of the same lesion over time. A lesion is defined to be of age 0 at the first timepoint of detection. Black numbers report the number of observations at the levels of leaf, lesion, and measurement interval. Total numbers for each wheat genotype are reported

Expressing lesion growth per unit of lesion perimeter yielded a measure that was relatively independent of lesion area at t_0_: While absolute lesion growth was strongly linearly correlated with lesion area at t_0_ (*R*^*2*^ = 0.35, *P* < 2.2 × 10^−16^), the correlation between perimeter-normalized lesion growth and lesion area at t_0_ was comparably weak (0.04 ≤ *R*^*2*^ ≤ 0.07, *P* < 2.2 × 10^−16^, Figure 4A, Supplementary Figure S10). This remaining relationship was monotonic and increasing but significantly non-linear, with growth levelling off for larger lesions. This indicated that larger lesions tended to grow faster, suggesting that their size reflected a higher growth rate rather than simply greater age. Initial lesion area (i.e., the area at the moment of its first detection) was also non-linearly associated with average lesion growth throughout the measurement period (*R*^*2*^ = 0.15, *P* < 2.2 × 10^−16^; Figure 4B). This general trend was observed for all host cultivars and across measurement batches and years. Lesion growth rate decreased exponentially with increasing lesion age (Figure 4C). Yet, after an initial rapid decay phase, lesions continued to grow at a relatively constant rate throughout the duration of measurements (Figure 4C). This finding confirmed frequent visual observations of continued lesion growth for extended periods of time without a clear end point. At first glance, these findings seem contradictory: larger lesions tended to grow faster, and older lesions tended to be larger, yet older lesions tended to grow more slowly. This apparent contradiction likely reflects the temporal dynamics of lesion development, where lesion growth initially accelerates with increasing size but eventually slows with age.

**Figure 4.**
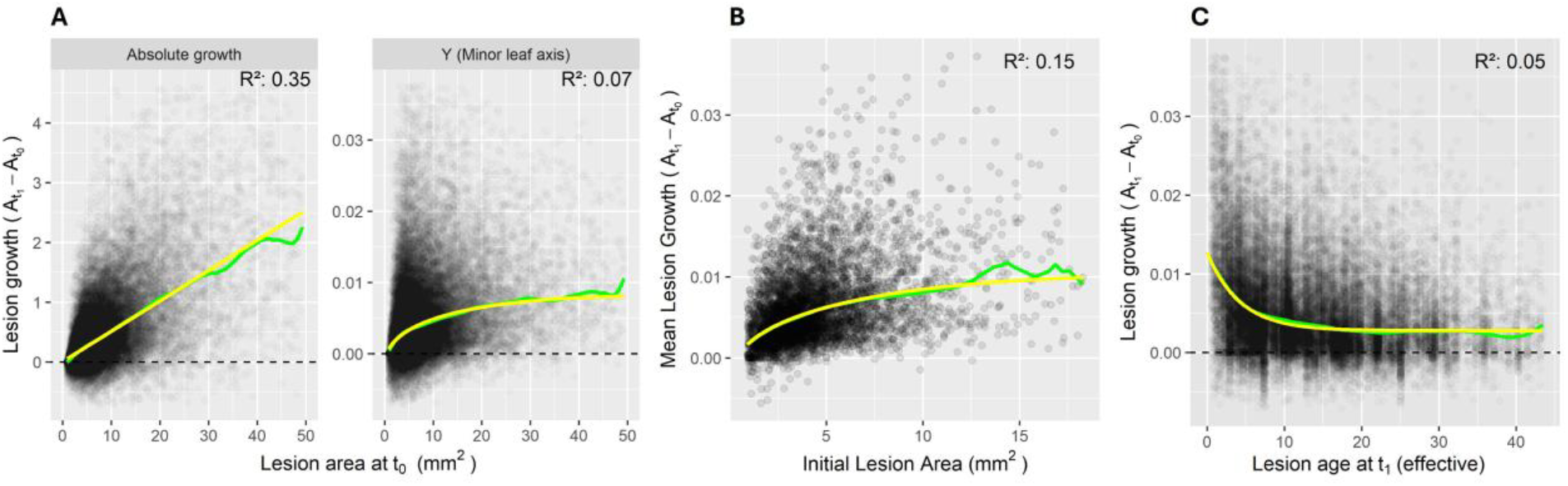
Lesion growth plotted against lesion area and lesion age. Yellow curves represent linear or non-linear quantile regression fits; green lines show loess-smoothed trends. X-axis limits were set to display all data points within the 99th percentile; models were fitted, and predictions are shown only within this range, due to limited data beyond this threshold. All coefficients of the regression models were statistically significant (p < 2.2 × 10^−16^). **(A)** Lesion growth vs. lesion area at the lag time point (t_0_). Growth is defined as the change in lesion area between consecutive time points 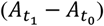, normalized by thermal time. It is plotted as absolute growth and per unit of lesion perimeter running along the minor (“Y”) leaf axes. **(B)** Mean lesion growth across all intervals vs. initial lesion size. Growth is defined here as change in lesion area between consecutive time points, normalized by thermal time and the length of the Y-perimeter of the lesion at t_0_. **(C)** Lesion growth against lesion age as measured in thermal time since its initial detection.

### Lesion growth prediction and influential features

RFE consistently identified environmental variables related to relative humidity and temperature during the growth interval as some of the most predictive features of lesion growth. Specifically, the mean and variance of relative humidity during the interval were the two most important features, with average ranks of 2.6 ± 1.07 and 1.8 ± 0.93, respectively (Figure 5B). Mean and variance of temperature were also consistently retained as informative predictors, with average ranks of 7.53 ± 1.91 and 4.77 ± 1.36, respectively. Other important features were related to the distance of the lesion perimeter from pycnidia and pycnidia density within the lesion as well as leaf and lesion age. Host cultivar (genotype_name) had an average rank of 10.2 ± 2.19. Year (exp_UID) and measurement batch representing the leaf layer and time in the growing season (batch_UID) were not amongst the most predictive features (Figure 5B).

**Figure 5.**
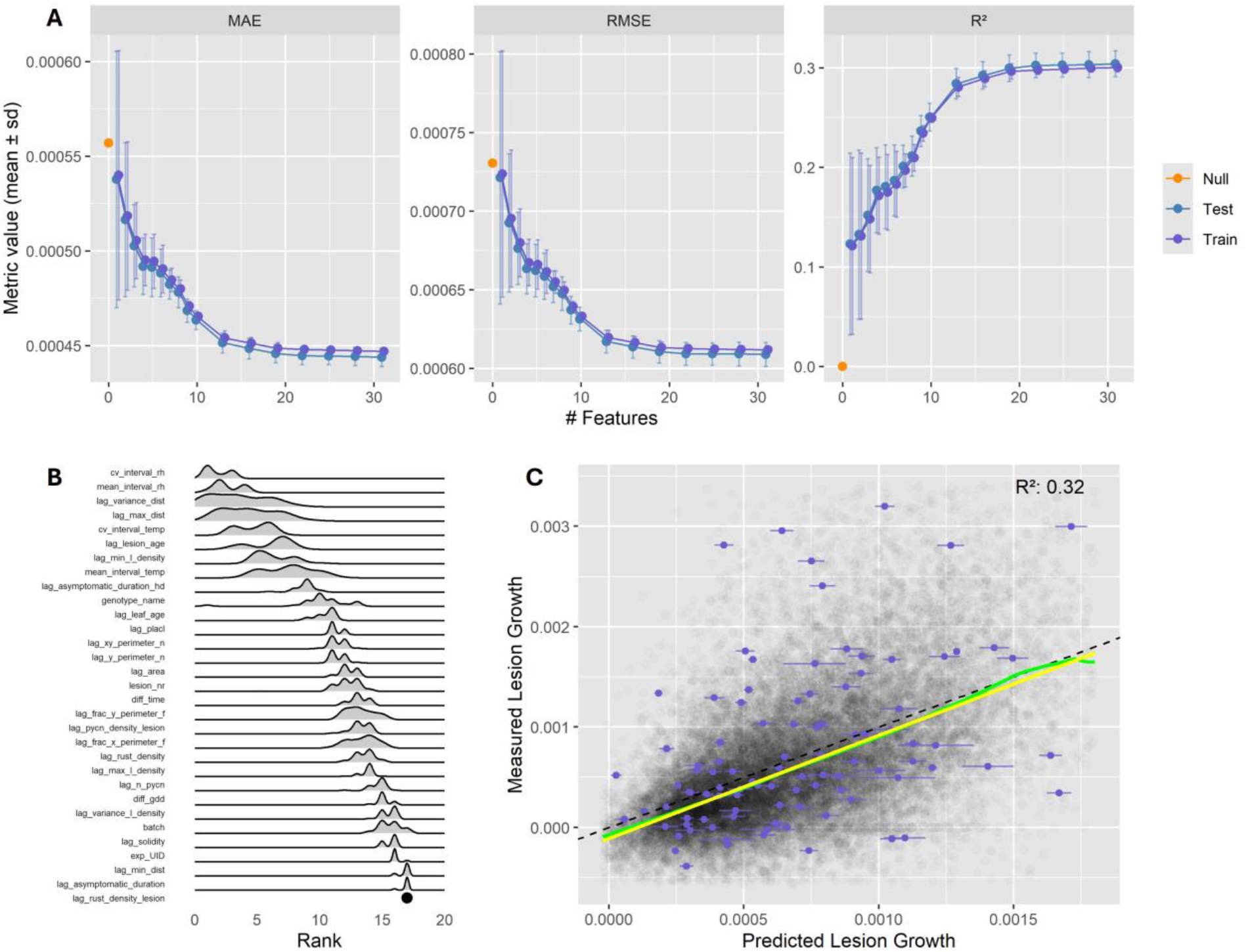
Prediction of lesion growth rates using disease phenomics features and environmental data. (**A**) Predictive model performance as a function of the number of features used during training, with means ± standard errors across 30 resamples. The ‘null’ performance is the mean of the response variable as baseline predictor. MAE – Mean average error; RMSE – Root mean squared error; R^2^ *-* coefficient of determination; (**B**) Distribution of importance ranks for the 20 most predictive features across 30 resamples. Low ranks indicate high relevance; the black dot represents invariant feature ranks across all resamples. (**C**) Measured versus predicted lesion growth using the 20 most predictive features. The yellow line is a linear quantile regression fit, the green line is a loess-smoothed trend, the dashed black line is the 1:1-line. Purple points with error bars are a small sub-sample showing means ± standard deviations of predicted values.

Optimal predictive performance was achieved when models were trained with around 20 of the most predictive features (Figure 5A). The average test RMSE for those models was 6.13 × 10^−4^. In comparison, using the mean of the response variable as a baseline predictor yielded a higher average RMSE of 7.31 × 10^−4^. The corresponding model trained on the full data set (i.e., on all available observations) reached an average cross-validated training RMSE of 6.06 × 10^−4^ and *R*^*2*^ of 0.32 (Figure 5C). In other words, approximately one third of the total variation in lesion growth could be explained based on the 20 most predictive features. Model performance varied substantially when only a few predictors were included but became more stable across resamples as the number of predictors increased (Figure 5A). Including additional features did not result in overfitting but instead led to small incremental improvements in model performance and contributed substantially to model stability. Notably, performance improved in a stepwise manner as additional features were included (Figure 5A). The high variability in model performance when only a few predictors were used, coupled with the fact that models with two predictors performed much better than models with a single predictor, indicates the importance of factor interactions where a feature becomes useful for prediction only once specific other features, providing more relevant context, are also included.

The direct relationship between lesion growth and environmental variables did not appear to follow a consistent overall trend (Supplementary Figure S11). While pairwise linear relationships between environmental covariates and lesion growth rate were statistically significant (*P* ≤ 8.7 × 10^−15^), the fraction of explained variance was very small (≤ 2%; Supplementary Figure S11). Strong confounding between environmental variables and experimental design factors such as batch and year (Figure 2B, C) as well as the variable performance of models trained using only relative humidity data (Figure 5A, B) suggested that the predictive power of these environmental covariates was overestimated and relied on modelling of local patterns that could often not be validated on test sets (cf. Supplementary Figure S11). This highlighted the need for complementary analyses that explicitly account for design structure, including year and batch effects.

Compared to environmental variables, pycnidiation patterns (maximum and variance of the distance of the lesion perimeter from nearest pycnidia, and minimum pycnidia density on the lesion perimeter) showed more consistent associations with lesion growth, explaining between 3% and 4% of variance in lesion growth (Supplementary Figure S12). These trends indicated that a greater distance of the lesion edge from pycnidia correlated with faster lesion growth. Several summary statistics of the distance between pycnidia and the lesion perimeter varied significantly and in a similar way with host cultivar (*F* = 123.5, *P* < 2.2 × 10^−16^) and, particularly, measurement batch (*F* = 645, *P* < 2.2 × 10^−16^; cf. Supplementary Figure S13). Thus, these features likely also captured a sizable portion of the variation in lesion growth rate associated with these factors, at least partly explaining their relatively high RFE ranks (Figure 5B).

High ranks of variables representing rust pustule density on the leaf (rust_density) and in the lesion (rust_density_lesion) indicated that co-infections on many leaves with brown rust, yellow rust, and STB did not affect STB lesion growth measurements (Figure 5B). This implied that rust occurrence either had no measurable impact on STB lesion growth or that potential interactions were effectively filtered out using diagnostic proxies derived from the image processing models during data pre-processing.

### Effects of host cultivar and environment on lesion growth

Wald tests for fixed effects in the mixed model confirmed the importance of all groups of factors considered (design-level, interval-level, and lesion-level). Lesion-level features (i.e., the maximum distance between pycnidia and the lesion perimeter, the area of the lesion, and lesion age) appeared to be most strongly associated with lesion growth and were all highly significant (Table 1). There was a highly significant batch effect (*P* < 2.2 × 10^−16^) whereas the year effect was not significant, suggesting limited influence of between-year environmental variation on lesion growth under the conditions of the study site. While mean interval temperature had a highly significant effect on lesion growth (*P* = 8.8 × 10^−12^), mean relative humidity did not, and this finding was somewhat unexpected given the results of feature selection (Figure 5B). However, note that interaction effects were considered in the random term and are analyzed separately.

**Table 1.**
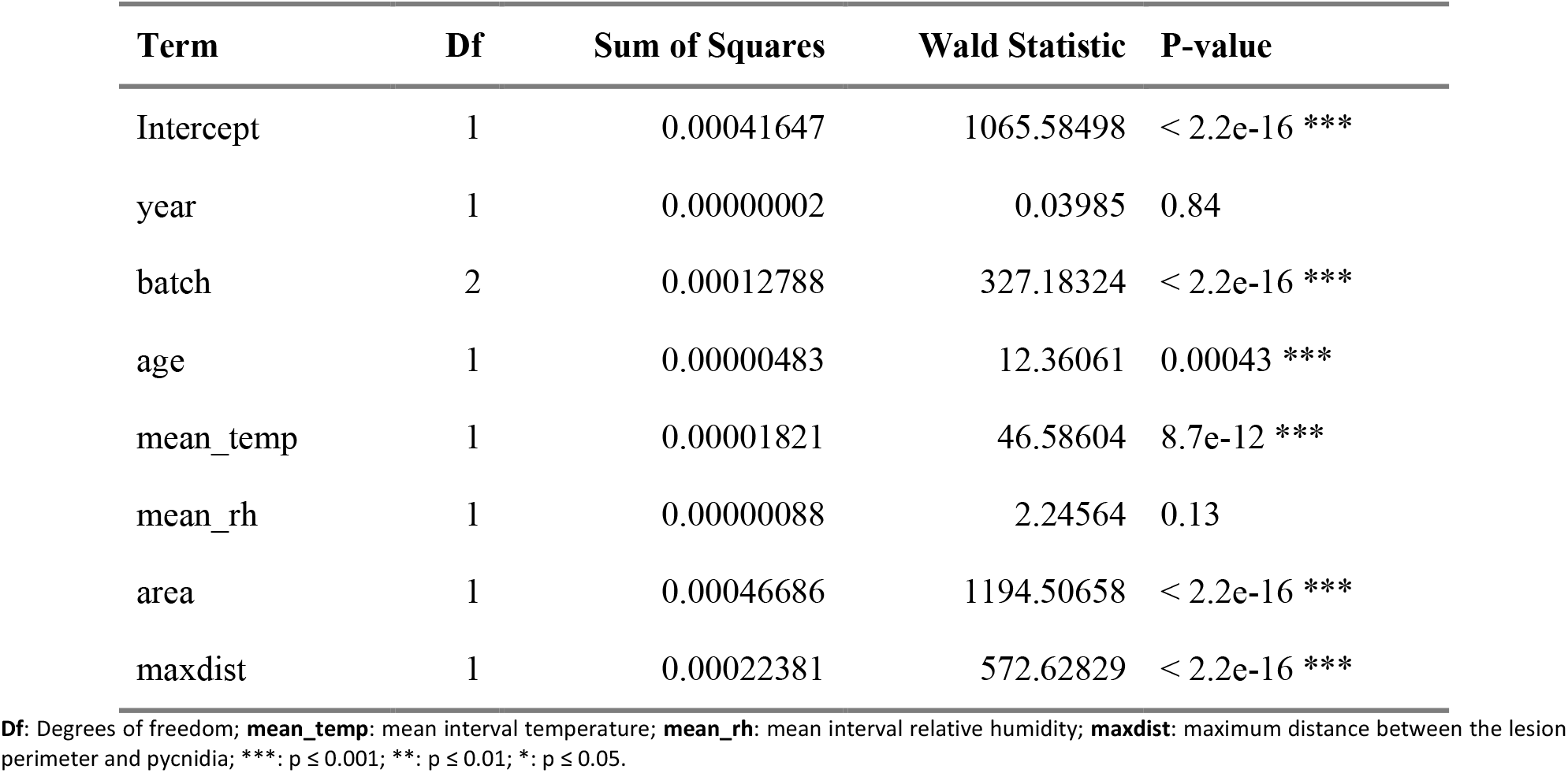
Wald tests for fixed effects in the linear mixed model analyzing lesion growth rate.

The proportion of the residual variation that was explained by the random effects (after adjusting for the fixed effects) was small, with 87.5% residual variance. Approximately 7% and 3.6% of the variance were explained by design factors (leaf and lesion), and only 1.8% was explained by host genotype. The proportion was even smaller for all interaction terms (< 1%) with genotype-by-temperature, genotype-by-humidity, and genotype-by-age interactions explaining increasingly less of the observed variance. Despite the low variance explained by these interaction terms, the heritability estimates for genotype-specific responses were moderate, with *h*^*2*^ = 0.46 for temperature, *h*^*2*^ = 0.40 for relative humidity, and *h*^*2*^ = 0.35 for lesion age. These are comparable to the heritability of the genotype main effect (*h*^*2*^ = 0.45), indicating that lesion growth in response to environmental conditions was clearly genotype-specific, even if the overall contribution of these interaction terms to the total variance was modest. For most genotypes, the estimated temperature response of lesion growth was substantial (Supplementary Figure S14).

### Relationship between lesion growth and overall QR

Within-time-point heritability of components of overall disease was very high (0.87 ≤ *h*^*2*^ ≤ 0.96 for disease incidence and 0.82 ≤ *h*^*2*^ ≤ 0.97 for conditional disease severity), except for conditional disease severity on one time point (t1 in 2023, *h*^*2*^ = 0.36). At this time point, only 3 replicate plots per cultivar under natural infection were evaluated, exhibiting generally low disease intensity. These results highlight the precision of the phenotyping for quantifying overall QR. Across time points, heritability remained high for disease incidence (*h*^*2*^ = 0.70) but dropped markedly for conditional severity (*h*^*2*^ = 0.12), likely reflecting interactions between genotype and leaf layer, growth stage, and environmental conditions. Confusion with physiological senescence at late time points cannot be excluded but should have been minimized by excluding these plots.

Genotypic BLUPs for disease incidence and conditional severity were highly correlated (*R*^*2*^ = 0.77, *P* = 1.87 × 10^−6^; Figure 6A). In contrast, lesion growth showed no correlation with either incidence or conditional severity across all 12 genotypes with available data (*R*^*2*^ ≤ 0.05, *P* ≥ 0.5; Figure 6B, 6C). However, cultivar ‘AUBUSSON’ represented a clear outlier (Cook’s Distance *D* = 3.51 vs. 0.08 ± 0.13 on average for the other observations). When cultivar ‘AUBUSSON’ was excluded, the linear correlation between lesion growth and conditional severity across the remaining 11 genotypes was strong and statistically significant (*R*^*2*^ = 0.58, *P* = 0.01; Figure 6B). The relationship with incidence remained not significant (*R*^*2*^ = 0.35, *P* = 0.06; Figure 6C).

**Figure 6.**
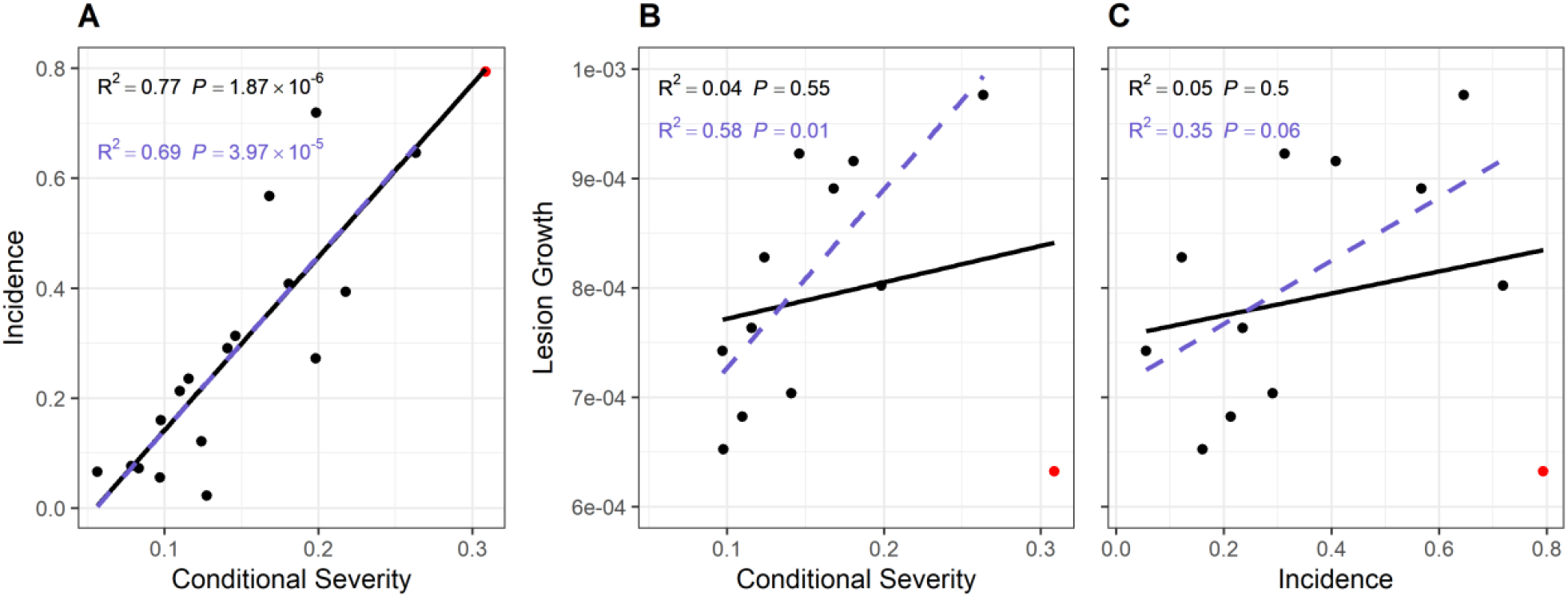
Components of quantitative resistance (QR) and their relationship with lesion growth. Genotypic BLUPs across two years are shown. Red points represent cultivar ‘AUBUSSON’. Black lines and text show linear regressions with R^2^ and *P*-values; purple lines and text show results after excluding ‘AUBUSSON’. (**A**) Disease incidence plotted against conditional severity; (**B**) Lesion growth plotted against conditional severity; (**C**) Lesion growth plotted against disease incidence.

Genotype-specific slopes describing the response of lesion growth to lesion age, relative humidity, and temperature showed no significant correlation with conditional severity when considering all genotypes (*R*^*2*^ ≤ 0.08, *P* ≥ 0.37; Supplementary Figure S15). Here, the presence of the outlier genotype ‘AUBUSSON’ had little effect on the direction of the associations. However, excluding this genotype suggested a significant negative correlation between the slope for relative humidity and conditional severity (*R*^*2*^ = 0.42, *P* = 0.03) and a non-significant trend toward a positive association with temperature (*R*^*2*^ = 0.27, *P* = 0.1); the effect of lesion age remained negligible (*R*^*2*^ = 0.09, *P* = 0.36).

These results suggested that the genotype main effect on lesion growth was more important for disease development than genotypic differences in the response of lesion growth to changes in environmental conditions. Main effect and responses to environmental conditions appeared to be relatively independent (*R*^*2*^ ≤ 0.13, *P* ≥ 0.2; Supplementary Figure S16), whereas the response to lesion age was significantly associated with the main effect (*R*^*2*^ = 0.28, *P* = 0.05; Supplementary Figure S16).

## Discussion

We employed a time-resolved image-based phenomics approach based on deep learning to quantify lesion growth on wheat leaves at scale and under naturally fluctuating environmental conditions, thereby capturing an important leaf-level component of seasonal STB epidemic development. To investigate the drivers of lesion growth, we combined predictive modelling with hypothesis-driven statistical modelling. This two-stage approach first enabled us to identify and rank key explanatory variables from diverse categories (lesion-level, leaf-level, measurement interval, and experimental factors) based on their predictive value. In the second stage, these insights guided the development of statistical models to estimate the specific contribution of host genotype and genotype-by-environment interactions as well as their heritability. Key findings from this study were that i) lesion growth rate was strongly associated with lesion-specific appearance traits which only partially aligned with host genotype, likely reflecting the combined influence of host background and the diversity of pathogen strains infecting each host; ii) the proportion of variance attributable to host genotype and genotype-by-environment interactions was relatively small; however, both main and some interaction effects were moderately heritable, indicating potential for selection in breeding for QR; and iii) lesion growth rate was not clearly associated with overall QR or with its individual components when all evaluated genotypes were considered. However, after excluding a single clear outlier genotype, a strong positive association with conditional severity emerged, implying that lesion growth may contribute substantially to overall field-level QR in most – but not all – wheat genotypes.

Few studies have investigated lesion growth rates under naturally fluctuating environmental conditions, or investigated their relationship to overall QR - and to the best of our knowledge, none have done so for STB. Most existing studies have relied on manual measurements of lesion size, which typically resulted in limited data sets covering a small number of cultivars and pathogen strains generated under highly controlled environment conditions. For STB, manual measurement of lesion size at the necessary scale is impractical due to the miniature size of symptoms and the rapid development and coalescence of lesions. We are aware of only one study comparable in scale to ours. In that study, SNB lesion growth rates were assessed across multiple cultivars and year-sites under naturally fluctuating environmental conditions and with genetically diverse pathogen populations (Adhikari *et al*., 2023). Although the authors did not directly estimate variance components, their results highlighted host genotype and temperature as key drivers of lesion growth: lesion growth rates were reported to be approximately four times higher in susceptible cultivars than in moderately resistant ones, and temperature had the strongest overall influence (Adhikari *et al*., 2023). Several other studies performed in various pathosystems under more controlled conditions also found strong effects of genotype (e.g., Sigulas *et al*., 1988; Koch & Mew, 1991; da Silva *et al*., 2012) and temperature (e.g., Pelletier & Fry, 1989; Paul & Munkvold, 2005) on lesion growth, and these effects explained a substantial portion of the total observed variation in lesion growth. Our study offers a contrasting view: Only a moderate portion (∼32%) of the overall variation in lesion growth rates was explainable (Figure 5C), and only a very small portion of the total variation could be attributed to host genotype and genotype-by-environment interactions. These contrasting results may be explained by several factors.

The similar study by Adhikari *et al*. (2023) addressed the wheat/*S. nodorum* pathosystem that shares many epidemiological processes with STB, but differs from the Wheat/*Z. tritici* pathosystem in several important aspects. *Z. tritici* has a clearly biphasic infection cycle characterized by a long latent period after infection during which host tissue colonization occurs without immediate symptom expression, followed by a necrotrophic phase that coincides with the development of necrotic symptoms (Sánchez-Vallet *et al*., 2015). At our study site, latent periods of three to four weeks are typically observed under field conditions when infections occur around flag leaf emergence. In contrast, *S. nodorum* has a much shorter latent period and a more necrotrophic behavior, often causing cell death while penetrating the leaf surface, and killing cells in advance of mycelial colonization (Friesen & Faris, 2021). The inverse gene-for-gene interactions characterizing SNB involve direct interactions between pathogen-secreted necrotrophic effectors and corresponding susceptibility factors in the host. These direct interactions could largely explain the strong genotype-specific differences in lesion development across host genotypes, although Adhikari *et al*. (2023) considered the involvement of other small-effect genes not directly involved in these interactions to be more likely. Nonetheless, compared to SNB, STB appears to be governed more by polygenic quantitative resistance mechanisms (Friesen & Faris, 2021) that would explain the smaller and less consistent genotype effects on STB lesion growth. Conversely, the highly quantitative nature of lesion growth within individual host genotypes supports the view that lesion expansion is governed by complex genetic interactions, involving both host and pathogen genotypes, with many pathogen genotypes often infecting the same leaf. The consistent association we observed between greater distances from pycnidia to the lesion perimeter and faster lesion growth (Figure 5A, Supplementary Figures S12, S13) could reflect inverse gene-for-gene interactions similar to those described in SNB, where necrosis can advance ahead of fungal colonization due to the effect of secreted necrotrophic effectors. This would corroborate a previously raised hypothesis based on the observation that the occurrence of chlorotic zones around necrotic lesions has a quantitative host-genetic basis (Anderegg *et al*., 2022).

By analyzing lesion growth from initial appearance to advanced age as a single process, we may have inadvertently confounded the dynamics of distinct biological processes that may differ in their response to genotype and environmental factors. The rapid early growth of lesions could reflect the transition from latent colonization to necrosis in the initially colonized leaf tissue, while slower growth observed at later stages would represent later invasion of new mycelium into surrounding, previously uncolonized tissue. Visual inspection provided some support for this hypothesis: for newly emerging lesions, often only a portion of a visible damaged zone was identified as a necrotic lesion, and this fraction rapidly increased across the first few observations (see e.g., lesion ‘34’ in Figure 1). Although a gradual decline in lesion growth rate over time has also been observed in SNB (Adhikari *et al*., 2023), the rapid exponential decay seen here (Figure 4C) may therefore not reflect aging of the lesion formed by the initial invasion of hyphae, but rather a shift between different phases of pathogenesis.

Besides the magnitude of host genotypic effects, the influence of environmental variables also contrasted strongly with previous findings from other pathosystems. For example, for early blight of potatoes, Pelletier & Fry (1989) found a very strong linear relationship between temperature and radial lesion growth when exposing leaves of a highly susceptible cultivar inoculated with a single field isolate to a wide range of constant temperatures (9-27°C). Similarly, Paul & Munkvold (2005) reported a significant effect of temperature on expansion of grey leaf spot lesions on maize leaves, especially when temperatures approached daily maximum temperatures normally occurring in the study region. Adhikari *et al*. (2023) reported a very strong influence of temperature on SNB lesion growth. Compared to most studies, the ranges in temperature and relative humidity observed within measurement batches in our study were narrow (Figure 2), and conditions may not have included clearly sub-optimal, stressful environments. While sequential measurement batches broadened the range of environmental conditions, this variation was not directly exploitable for modelling the genotype-by-environment interactions, since we included a fixed batch effect. In contrast, Adhikari *et al*. (2023) stated that “*Cohort* [here: batch] *was not included in the model because it was inevitably confounded with temperature and relative humidity*.” This was also evident in our data (Figure 2). However, studies in other pathosystems provided strong evidence that lesion growth measurements are significantly influenced by leaf or plant age and phenology, as well as by the seasonal timing of measurements (e.g., Emge *et al*., 1975; Pelletier & Fry, 1989). We therefore chose to include a batch effect to control for systematic differences between measurements taken at different phenological stages of the crop, including different leaf layers, and under broadly contrasting environmental conditions, including diurnal variation, that are not fully captured by mean temperature and mean relative humidity alone. Including batch effects also helped mitigate the significant imbalance in the data set arising from the strictly incidence-based selection of experimental plots. We consider this a more cautious approach that helps control for unknown and unmodeled sources of variation across sequential measurement batches, yielding more conservative but also more robust estimates of genotype and genotype-by-environment interaction effects.

Some studies (Bernard *et al*., 2013, Bernard *et al*., 2022) pointed out that air temperature recorded at a nearby weather station may be inappropriate to study the effect of temperature on pathogen development in infected leaves due to discrepancies between ambient air and actual leaf temperature. Here, our focus was on a relative comparison of host cultivars rather than a characterization of the absolute temperature response of pathogen development. Therefore, consistent discrepancies between leaf and air temperature as analyzed by Bernard *et al*. (2022) are less problematic, provided they affect all cultivars similarly. We recognize that a portion of the discrepancy is likely to be genotype-specific, for example due to differences in leaf orientation and consequent exposure to direct sunlight during the day (Anderegg *et al*., 2021). Such differences may shift the mean leaf temperature across genotypes and increase within-leaf or within-canopy temperature variation. In applied field settings, capturing this fine-scale temperature heterogeneity and mapping it to lesion growth patterns is impractical. As a result, any genotype-specific deviations in actual leaf temperature will be confounded with genuine differences in temperature sensitivity of lesion growth. The same will be true for relative humidity within canopies, which can vary as a function of canopy three-dimensional structure and density. While the impracticality of disentangling this complexity under field conditions represents a limitation, the overall outcomes still reflect realistic, integrated genotype responses under field conditions which are relevant for evaluating QR in applied breeding scenarios.

The moderate heritability of both genotype and genotype-by-environment interaction effects on lesion growth, despite their small variance contributions, suggest that selection for these traits is possible but may require high precision and replication. Compared to many other foliar diseases (e.g., northern leaf blight on corn; Sigulas *et al*., 1988), a very high precision is needed to reliably measure growth of STB lesions over short time periods. A high measurement frequency was required to capture the highly dynamic growth process and to avoid treating separate nearby infections as single lesions. Longer measurement intervals (e.g., 2-to 5-day intervals for SNB monitoring; Adhikari *et al*., 2023) would have resulted in substantial confounding of lesion growth and lesion coalescence and would not have yielded a satisfactory number of repeat assessments for most tracked lesions. While we believe that our current method offers higher precision than manual assessments, further improvements are certainly possible. In particular, enhancing the accuracy of image alignment and lesion segmentation in the image time series—potentially by addressing both tasks simultaneously—could further increase the reliability and resolution of lesion growth measurements. The large, manually curated data set underlying this work can serve as a solid basis for the development of such methods.

Our approach to measuring lesion growth rates accounted for lesion shape and spatial context to some extent, for example by relating lesion growth to lesion perimeter length and normalizing for the presence of obstacles to growth. Yet, we treated lesion growth as an integrated lesion-level metric rather than modelling it in a spatially explicit manner. In contrast, Leclerc *et al*. (2023) proposed the use of a reaction-diffusion framework to explicitly model the growth of individual lesions as a continuous function of space and time. While this offers a promising approach, the data analyzed herein represent a considerably more complex system than theirs in terms of lesion spatial context, irregular growth patterns, fluctuating environmental conditions, and the diversity of host-pathogen-interactions. Substantial extensions of the proposed modelling framework will therefore be required to accommodate this increased complexity.

The absence of a significant correlation between lesion growth rate and overall QR (Figure 6B, 6C) may suggest that lesion growth played a limited role in shaping STB epidemics under the conditions of our experiment. However, excluding a clear outlier host genotype identified already in the first year (Anderegg *et al*., 2024b) and confirmed in the second, resulted in a different picture, with a strong and statistically significant positive correlation emerging between lesion growth and conditional severity among the remaining genotypes (Figure 6B). It is important to note that the outlier genotype appeared to represent a true biological outlier since no technical issues related to lesion growth measurements could be identified and trait estimates were based on a substantial number of observations (Figure 3). While slow lesion growth suggested that this cultivar harbors some form of QR, at the same time it appeared to be highly susceptible to new infections, to an extent that nullified the potentially beneficial effect of reduced lesion growth on overall QR (Anderegg *et al*., 2024b). This highlights how different components of rate-reducing QR may not necessarily be correlated, a finding already reported for other pathosystems under more controlled conditions (Jeger *et al*., 1983; Loughman *et al*., 1996). Determining the overall contribution of lesion growth to field epidemics of STB may therefore require broader studies involving a larger set of genotypes and ideally a more complete characterization of QR components.

By combining deep-learning-based phenomics with advanced statistical modelling, we achieved the most precise and large-scale quantification of STB lesion growth under field conditions. These unique observations revealed the complexity of lesion development under natural epidemic conditions and highlighted the challenge of using lesion growth rates as a standalone measure of QR. The high precision and the statistical power stemming from the large number of observations enabled the detection of heritable genetic effects on lesion growth, facilitating the identification of distinct types of QR. Thus, our approach paves the way for a more complete functional understanding of resistance mechanisms and a more targeted use of these components of QR in breeding for durable resistance.

## Acknowledgements

We thank Stefanie Dräyer, Seraina Wagner, Mathilda Pohier, Yurun Jin, Lisa Hohl, Tenzin Khampo, Vilma Rantanen, and Giulia Malingamba (all ETH Plant Pathology) for their contributions to field data collection and image annotation; Andreas Hund and Achim Walter (both ETH Crop Science) for sharing field and computing resources; and Simon Corrado and Brigitta Herzog (both ETH Crop Science) for their support in crop husbandry, experiment planning, and seed preparation and management. The project was funded by ETH Zürich.

## Author contributions

JA conceptualized the study, planned and coordinated the measurement campaigns, and curated the data. JA and RZ conducted field experiments. JA and LR analyzed the data with input from BAM. RZ supported image processing. BAM acquired project funding. JA wrote the original draft of the manuscript. BAM and LR reviewed and edited the manuscript.

## Data availability

Data acquired and analyzed in this study are available via the Repository for Publications and Research data of ETH Zürich (https://doi.org/10.3929/ethz-b-000735494). All code required to reproduce the numerical results, figures, and tables presented in the manuscript is publicly available at https://github.com/and-jonas/septoria-lesion-growth. Raw image data will be made available by the authors upon reasonable request, without undue reservation. A sample data set is available at https://doi.org/10.3929/ethz-b-000735497. Code required to process raw image data and extract lesion-level data as analyzed in this paper is publicly available at https://github.com/and-jonas/sympathique-wheat/tree/development. Stable releases will follow upon acceptance of the manuscript.

## References

Adhikari U, Brown J, Ojiambo PS, Cowger C. 2023. Effects of Host and Weather Factors on the Growth Rate of Septoria nodorum Blotch Lesions on Winter Wheat. Phytopathology® 113: 1898–1907.

Alassimone J, Praz C, Lorrain C, De Francesco A, Carrasco-López C, Faino L, Shen Z, Meile L, Sánchez-Vallet A. 2024. The Zymoseptoria tritici Avirulence Factor AvrStb6 Accumulates in Hyphae Close to Stomata and Triggers a Wheat Defense Response Hindering Fungal Penetration. Molecular Plant-Microbe Interactions® 37: 432–444.

Anderegg J, Aasen H, Perich G, Roth L, Walter A, Hund A. 2021. Temporal trends in canopy temperature and greenness are potential indicators of late-season drought avoidance and functional stay-green in wheat. Field Crops Research 274: 108311.

Anderegg J, Hund A, Karisto P, Mikaberidze A. 2019. In-Field Detection and Quantification of Septoria Tritici Blotch in Diverse Wheat Germplasm Using Spectral–Temporal Features. Frontiers in Plant Science 10.

Anderegg J, Kirchgessner N, Aasen H, Zumsteg O, Keller B, Zenkl R, Walter A, Hund A. 2024a. Thermal imaging can reveal variation in stay-green functionality of wheat canopies under temperate conditions. Frontiers in Plant Science 15.

Anderegg J, Kirchgessner N, Kronenberg L, McDonald BA. 2022. Automated Quantitative Measurement of Yellow Halos Suggests Activity of Necrotrophic Effectors in Septoria tritici Blotch. Phytopathology® 112: 2560–2573.

Anderegg J, Yu K, Aasen H, Walter A, Liebisch F, Hund A. 2020. Spectral Vegetation Indices to Track Senescence Dynamics in Diverse Wheat Germplasm. Frontiers in Plant Science 10.

Anderegg J, Zenkl R, Kirchgessner N, Hund A, Walter A, McDonald BA. 2024b. SYMPATHIQUE: image-based tracking of symptoms and monitoring of pathogenesis to decompose quantitative disease resistance in the field. Plant Methods 20: 170.

Anderegg J, Zenkl R, Walter A, Hund A, McDonald BA. 2023. Combining High-Resolution Imaging, Deep Learning, and Dynamic Modeling to Separate Disease and Senescence in Wheat Canopies. Plant Phenomics 5: 0053.

Armour T, Viljanen-Rollinson SLH, Chng SF, Butler RC, Jamieson PD, Zyskowski RF. 2004. Examining the latent period of Septoria tritici blotch in a field trial of winter wheat. New Zealand Plant Protection 57: 116–120.

Battache M, Lebrun M-H, Sakai K, Soudière O, Cambon F, Langin T, Saintenac C. 2022. Blocked at the Stomatal Gate, a Key Step of Wheat Stb16q-Mediated Resistance to Zymoseptoria tritici. Frontiers in Plant Science 13.

Battache M, Suarez-Fernandez M, Klooster MV, Cambon F, Sánchez-Vallet A, Lebrun M-H, Langin T, Saintenac C. 2024. Stomatal penetration: the cornerstone of plant resistance to the fungal pathogen Zymoseptoria tritici. BMC Plant Biology 24: 736.

Behmann J, Bohnenkamp D, Paulus S, Mahlein A-K. 2018. Spatial Referencing of Hyperspectral Images for Tracing of Plant Disease Symptoms. Journal of Imaging 4: 143.

Berger RD, Filho AB, Amorim L. 1997. Lesion Expansion as an Epidemic Component. Phytopathology® 87: 1005–1013.

Bernard F, Chelle M, Fortineau A, Riahi El Kamel O, Pincebourde S, Sache I, Suffert F. 2022. Daily fluctuations in leaf temperature modulate the development of a foliar pathogen. Agricultural and Forest Meteorology 322: 109031.

Bernard F, Sache I, Suffert F, Chelle M. 2013. The development of a foliar fungal pathogen does react to leaf temperature! New Phytologist 198: 232–240.

Brokenshire T. 1976. The reaction of wheat genotypes to Septoria tritici. Annals of Applied Biology 82: 415–423.

Brown JKM, Chartrain L, Lasserre-Zuber P, Saintenac C. 2015. Genetics of resistance to Zymoseptoria tritici and applications to wheat breeding. Fungal Genetics and Biology 79: 33–41.

Butler DG, Cullis BR, Gilmour AR, Gogel BJ, Thompson R. 2018. ASReml estimates variance components under a general linear mixed model by residual maximum likelihood (REML). R Package version 4.2.0.355. : 188.

Chaloner TM, Fones HN, Varma V, Bebber DP, Gurr SJ. 2019. A new mechanistic model of weather-dependent Septoria tritici blotch disease risk. Philosophical Transactions of the Royal Society B: Biological Sciences 374: 20180266.

Coombes N. 2009. DiGGer. Design search tool in R.

Cowger C, Brown JKM. 2019. Durability of Quantitative Resistance in Crops: Greater Than We Know? Annual Review of Phytopathology 57: 253–277.

Cullis BR, Smith AB, Coombes NE. 2006. On the design of early generation variety trials with correlated data. Journal of Agricultural, Biological, and Environmental Statistics 11: 381–393.

Díaz-Lago JE, Stuthman DD, Leonard KJ. 2003. Evaluation of Components of Partial Resistance to Oat Crown Rust Using Digital Image Analysis. Plant Disease 87: 667–674.

Emge RG, Kingsolver CH, Johnson DR. 1975. Growth of the Sporulating Zone of Puccinia striiformis and Its Relationship to Stripe Rust Epiphytology. Phytopathology 65: 679.

Ficke A, Cowger C, Bergstrom G, Brodal G. 2018. Understanding Yield Loss and Pathogen Biology to Improve Disease Management: Septoria Nodorum Blotch - A Case Study in Wheat. Plant Disease 102: 696–707.

Friesen TL, Faris JD. 2021. Characterization of Effector–Target Interactions in Necrotrophic Pathosystems Reveals Trends and Variation in Host Manipulation. Annual Review of Phytopathology 59: 77–98.

Guyon I, Weston J, Barnhill S, Vapnik V. 2002. Gene Selection for Cancer Classification using Support Vector Machines. Machine Learning 46: 389–422.

Jeger MJ, Gareth Jones D, Griffiths E. 1983. Components of partial resistance of wheat seedlings to Septoria nodorum. Euphytica 32: 575–584.

Jørgensen LN, Olsen LV. 2007. Control of tan spot (Drechslera tritici-repentis) using cultivar resistance, tillage methods and fungicides. Crop Protection 26: 1606–1616.

Karisto P, Dora S, Mikaberidze A. 2019. Measurement of infection efficiency of a major wheat pathogen using time-resolved imaging of disease progress. Plant Pathology 68: 163–172.

Karisto P, Hund A, Yu K, Anderegg J, Walter A, Mascher F, McDonald BA, Mikaberidze A. 2018. Ranking Quantitative Resistance to Septoria tritici Blotch in Elite Wheat Cultivars Using Automated Image Analysis. Phytopathology 108: 568–581.

Karisto P, Suffert F, Mikaberidze A. 2022. Measuring Splash Dispersal of a Major Wheat Pathogen in the Field. PhytoFrontiersTM 2: 30–40.

Koch MF, Mew TW. 1991. Rate of Lesion Expansion in Leaves as a Parameter of Resistance to Xanthomonas campestris pv. oryzae in Rice. Plant Disease 75: 897.

Koenker R, code) SP (Contributions to CQ, code) PTN (Contributions to SQ, code) BM (Contributions to preprocessing, code) AZ (Contributions to dynrq code essentially identical to his dynlm, code) PG (Contributions to nlrq, routines) CM (author of several linpack, sparskit2) YS (author of, code) VC (contributions to extreme value inference, code) IF-V (contributions to extreme value inference, et al. 2025. quantreg: Quantile Regression.

Koenker R, Machado JAF. 1999. Goodness of Fit and Related Inference Processes for Quantile Regression. Journal of the American Statistical Association 94: 1296–1310.

Kuhn M, Wing J, Weston S, Williams A, Keefer C, Engelhardt A, Cooper T, Mayer Z, Kenkel B, R Core Team, et al. 2021. caret: Classification and Regression Training.

Lancashire PD, Jones DGareth. 1985. Components of partial resistance to Septoria nodorum in winter wheat. Annals of Applied Biology 106: 541–553.

Langlands-Perry C, Pitarch A, Lapalu N, Cuenin M, Bergez C, Noly A, Amezrou R, Gélisse S, Barrachina C, Parrinello H, et al. 2023. Quantitative and qualitative plant-pathogen interactions call upon similar pathogenicity genes with a spectrum of effects. Frontiers in Plant Science 14.

Leclerc M, Jumel S, Hamelin FM, Treilhaud R, Parisey N, Mammeri Y. 2023. Imaging with spatio-temporal modelling to characterize the dynamics of plant-pathogen lesions. PLOS Computational Biology 19: e1011627.

Lovell DJ, Hunter T, Powers SJ, Parker SR, Van den Bosch F. 2004. Effect of temperature on latent period of septoria leaf blotch on winter wheat under outdoor conditions. Plant Pathology 53: 170–181.

Magboul AM, Geng S, Gilchrist DG, Jackson LF. 1992. Environmental Influence on the Infection of Wheat by Mycosphaerella graminicola. Phytopathology 82: 1407.

McDonald BA, Suffert F, Bernasconi A, Mikaberidze A. 2022. How large and diverse are field populations of fungal plant pathogens? The case of Zymoseptoria tritici. Evolutionary Applications 15: 1360–1373.

Meile L, Croll D, Brunner PC, Plissonneau C, Hartmann FE, McDonald BA, Sánchez-Vallet A. 2018. A fungal avirulence factor encoded in a highly plastic genomic region triggers partial resistance to septoria tritici blotch. New Phytologist 219: 1048–1061.

Meile L, Garrido-Arandia M, Bernasconi Z, Peter J, Schneller A, Bernasconi A, Alassimone J, McDonald BA, Sánchez-Vallet A. 2023. Natural variation in Avr3D1 from Zymoseptoria sp. contributes to quantitative gene-for-gene resistance and to host specificity. New Phytologist 238: 1562–1577.

Parlevliet JE. 1975. Partial resistance of barley to leafrust, Puccinia hordei. I. Effect of cultivar and development stage on latent period. Euphytica 24: 21–27.

Parlevliet JE. 1979. Components of Resistance that Reduce the Rate of Epidemic Development. Annual Review of Phytopathology 17: 203–222.

Paul PA, Munkvold GP. 2005. Influence of Temperature and Relative Humidity on Sporulation of Cercospora zeae-maydis and Expansion of Gray Leaf Spot Lesions on Maize Leaves. Plant Disease 89: 624–630.

Pelletier JR Fry. 1989. Characterization of Resistance to Early Blight in Three Potato Cultivars: Incubation Period, Lesion Expansion Rate, and Spore Production. Phytopathology 79: 511.

Piepho H-P, Möhring J, Schulz-Streeck T, Ogutu JO. 2012. A stage-wise approach for the analysis of multi-environment trials. Biometrical Journal 54: 844–860.

Poland JA, Balint-Kurti PJ, Wisser RJ, Pratt RC, Nelson RJ. 2009. Shades of gray: the world of quantitative disease resistance. Trends in Plant Science 14: 21–29.

Rodriguez-Alvarez MX, Lee D-J, Kneib T, Durban M, Eilers P. 2019. SAP: Multidimensional Generalized P-Splines Regression Models Estimation.

Roth L, Binder M, Kirchgessner N, Tschurr F, Yates S, Hund A, Kronenberg L, Walter A. 2024. From Neglecting to Including Cultivar-Specific Per Se Temperature Responses: Extending the Concept of Thermal Time in Field Crops. Plant Phenomics 6: 0185.

Roth L, Rodríguez-Álvarez MX, van Eeuwijk F, Piepho H-P, Hund A. 2021. Phenomics data processing: A plot-level model for repeated measurements to extract the timing of key stages and quantities at defined time points. Field Crops Research 274: 108314.

Sánchez-Vallet A, McDonald MC, Solomon PS, McDonald BA. 2015. Is Zymoseptoria tritici a hemibiotroph? Fungal Genetics and Biology 79: 29–32.

Savary S, Willocquet L, Pethybridge SJ, Esker P, McRoberts N, Nelson A. 2019. The global burden of pathogens and pests on major food crops. Nature Ecology & Evolution 3: 430–439.

Schilly A, Risser P, Ebmeyer E, Hartl L, Reif JC, Würschum T, Miedaner T. 2011. Stability of Adult-plant Resistance to Septoria tritici blotch in 24 European Winter Wheat Varieties Across Nine Field Environments. Journal of Phytopathology 159: 411–416.

Shaner G. 1973. Evaluation of Slow-Mildewing Resistance of Knox Wheat in the Field. Phytopathology 63: 867.

Sigulas KM, Hill RR, Ayers JE. 1988. Genetic Analysis of Exserohilum turcicum Lesion Expansion on Corn. Phytopathology 78: 149.

da Silva MR, Martinelli JA, Federizzi LC, Chaves MS, Pacheco MT. 2012. Lesion size as a criterion for screening oat genotypes for resistance to leaf spot. European Journal of Plant Pathology 134: 315–327.

Simon MR, Cordo CA. 1997. Inheritance of partial resistance to Septoria tritici in wheat (Triticum aestivum): limitation of pycnidia and spore production. Agronomie 17: 343–347.

Simon MR, Cordo CA. 1998. Diallel analysis of four resistance components to Septoria tritici in six crosses of wheat (Triticum aestivum). Plant Breeding 117: 123–126.

Stewart EL, Hagerty CH, Mikaberidze A, Mundt CC, Zhong Z, McDonald BA. 2016. An Improved Method for Measuring Quantitative Resistance to the Wheat Pathogen Zymoseptoria tritici Using High-Throughput Automated Image Analysis. Phytopathology 106: 782–788.

Suffert F, Sache I. 2011. Relative importance of different types of inoculum to the establishment of Mycosphaerella graminicola in wheat crops in north-west Europe. Plant Pathology 60: 878–889.

Willocquet L, Savary S, Yuen J. 2017. Multiscale Phenotyping and Decision Strategies in Breeding for Resistance. Trends in Plant Science 22: 420–432.

Wójtowicz A, Wójtowicz, Marek, Sigvald, Roland, and Pasternak M. 2017. Predicting the effects of climate change on the latency period of wheat leaf rust in western Poland. Acta Agriculturae Scandinavica, Section B — Soil & Plant Science 67: 223–234.

Wright MN, Ziegler A. 2017. ranger: A Fast Implementation of Random Forests for High Dimensional Data in C++ and R. Journal of Statistical Software 77.

Zenkl R, McDonald BA, Walter A, Anderegg J. 2025. Towards high throughput in-field detection and quantification of wheat foliar diseases using deep learning. Computers and Electronics in Agriculture 232: 109854.

