## Supplementary Materials for "Tracking Lesion Growth in the Field: Imaging and Deep Learning Reveal Components of Quantitative Resistance"

### Supplementary Figures

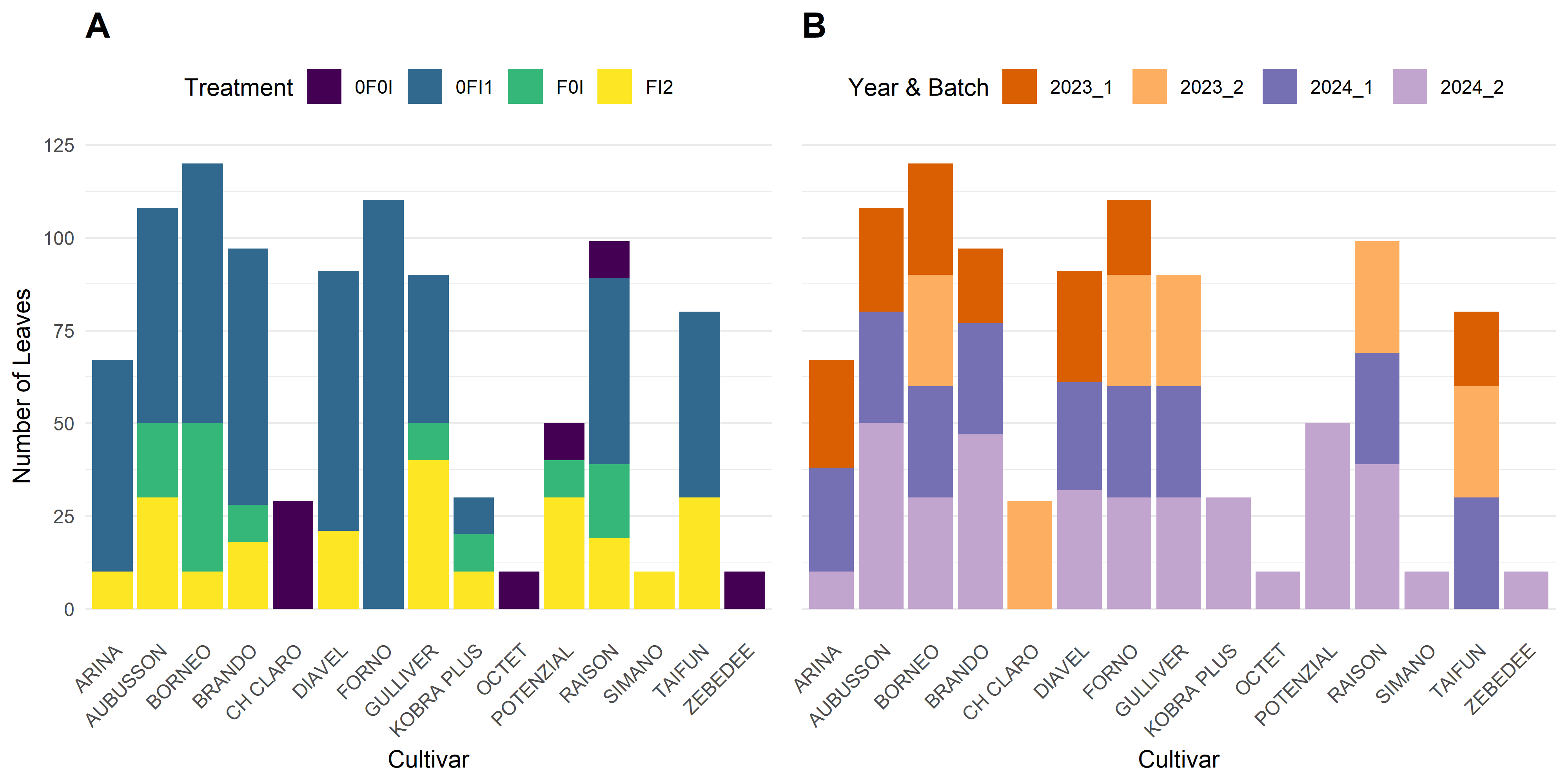

**Supplementary Figure S 1** Distribution of sampled leaves using different criteria. (**A**) Sample distribution by cultivar and treatment for pooled data from both years. (**B**) Sample distribution by year and measurement batch, for pooled data from all treatments. Batch 1 corresponds to the penultimate leaves that were monitored first; Batch 2 corresponds to the flag leaves that were monitored later.

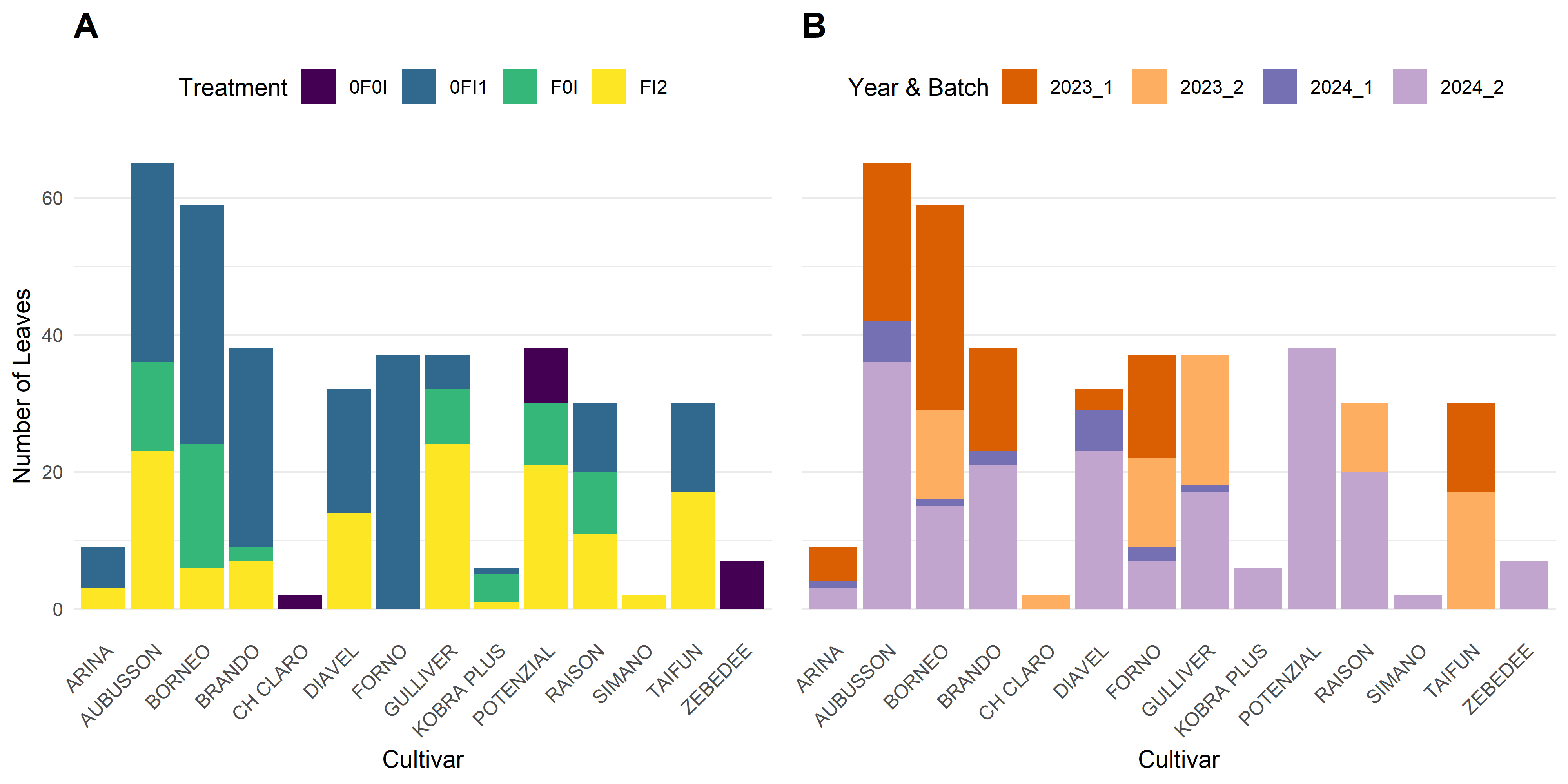

**Supplementary Figure S 2** Distribution of sampled leaves that were retained for the analysis, using different criteria. (**A**) Sample distribution by cultivar and treatment for pooled data from both years. (**B**) Sample distribution by year and measurement batch, for pooled data from all treatments. Batch 1 corresponds to the penultimate leaves that were monitored first; Batch 2 corresponds to the flag leaves that were monitored later.

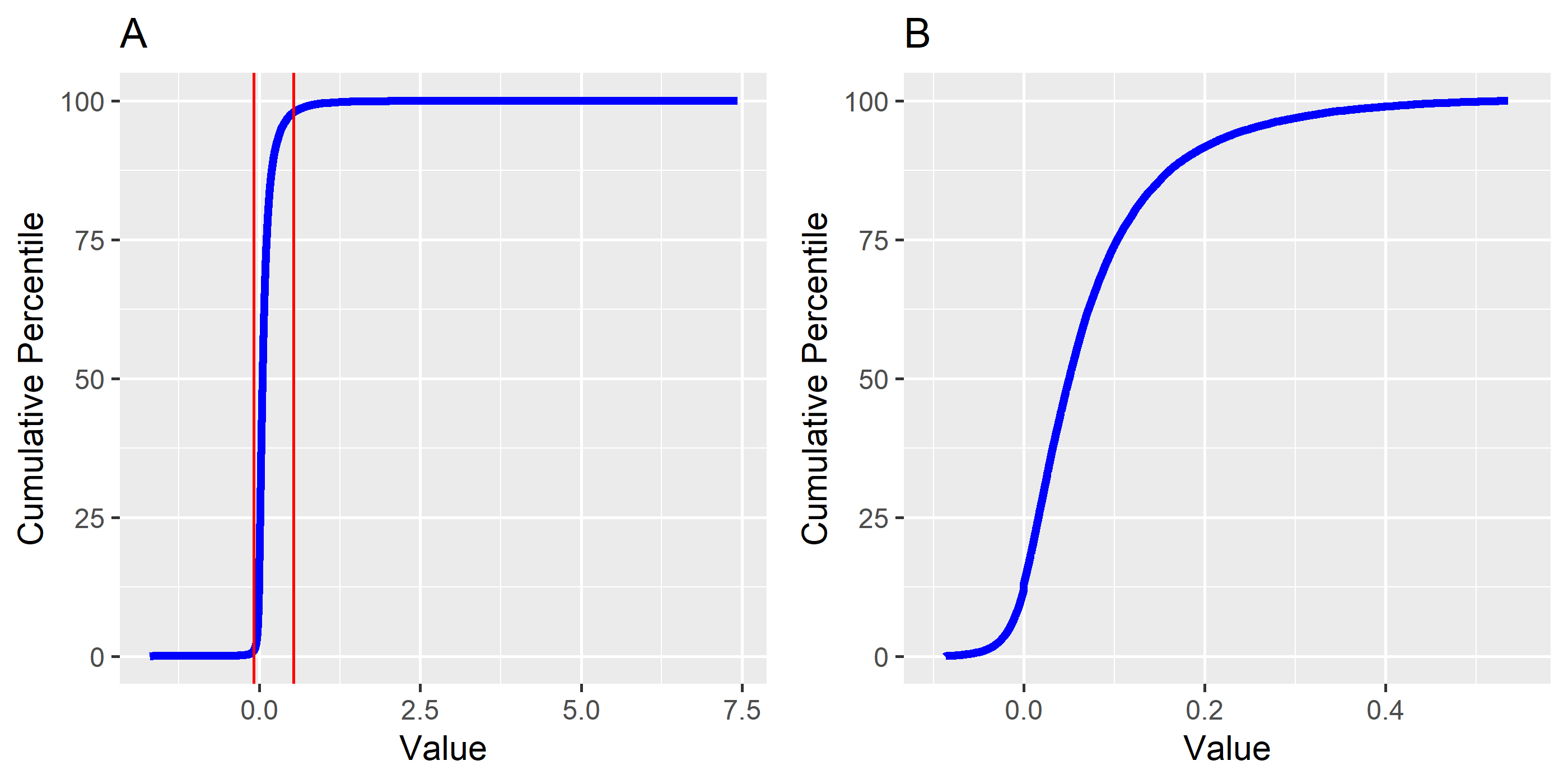

**Supplementary Figure S 3** Cumulative percentile distribution of lesion growth data before (**A**) and after trimming of outlier observations (**B**). The distribution in (A) reveals many extreme observations on both ends of the distribution. Red lines in (A) mark the applied thresholds corresponding to the 1^st^ and 98^th^ percentiles.

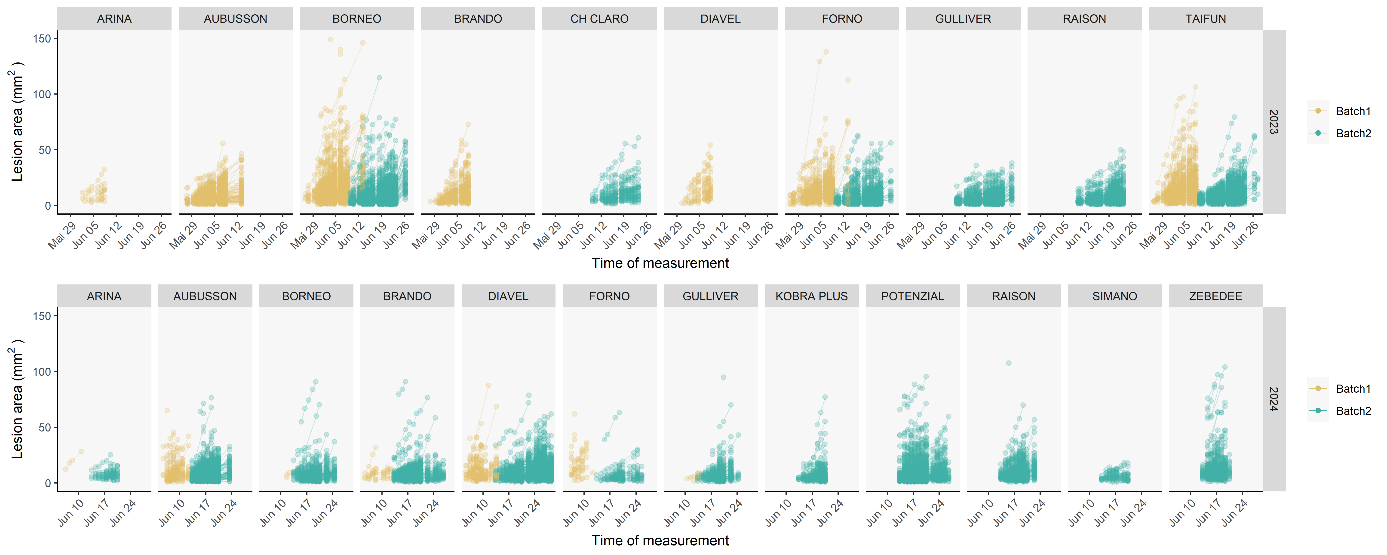

**Supplementary Figure S 4** Lesion area plotted against timepoint of measurement, shown separately for each wheat cultivar in each year. Lines connect repeat measurements of the same lesion over time. New lesions continue to appear throughout the duration of the measurement.

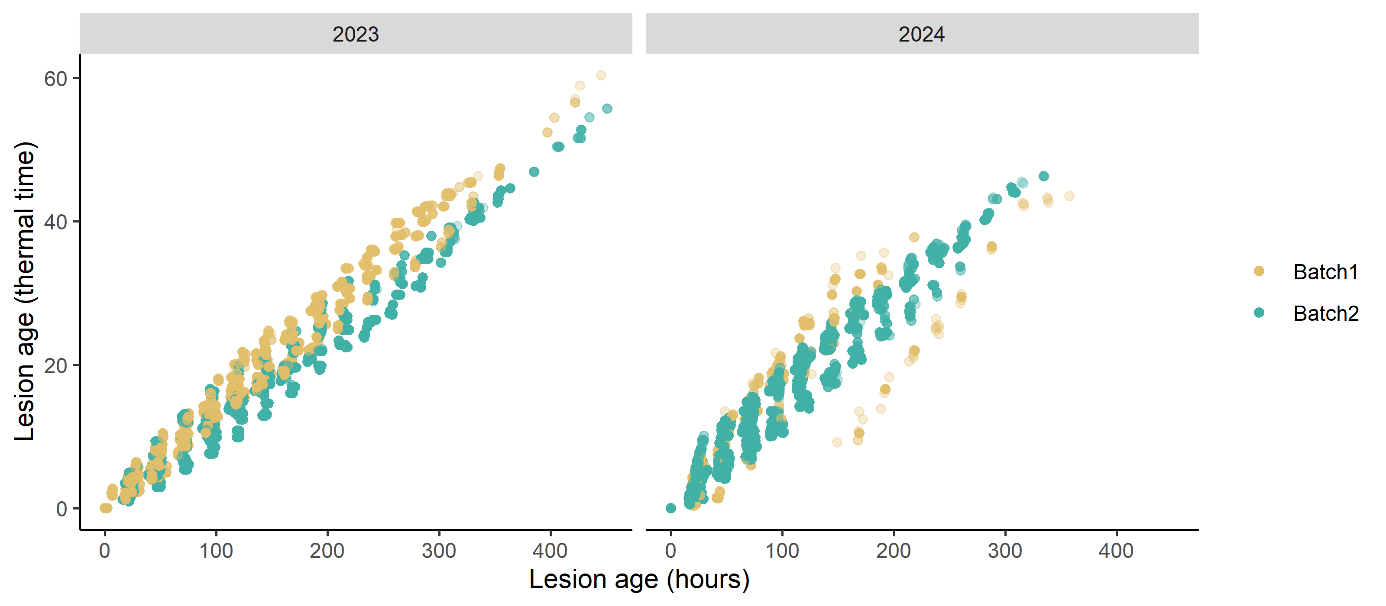

**Supplementary Figure S 5** Relationship between lesion age in terms of chronological and effective thermal (temperature-corrected) time across the years and including the two measurement batches in each experiment (Batch1 – penultimate leaves, and Batch2 – flag leaves). At constant optimal temperatures for lesion growth (estimated at 23°C; Chaloner et al. 2019), chronological time in hours is equivalent to effective thermal time. Reduced effective thermal times indicate suboptimal temperatures for lesion growth.

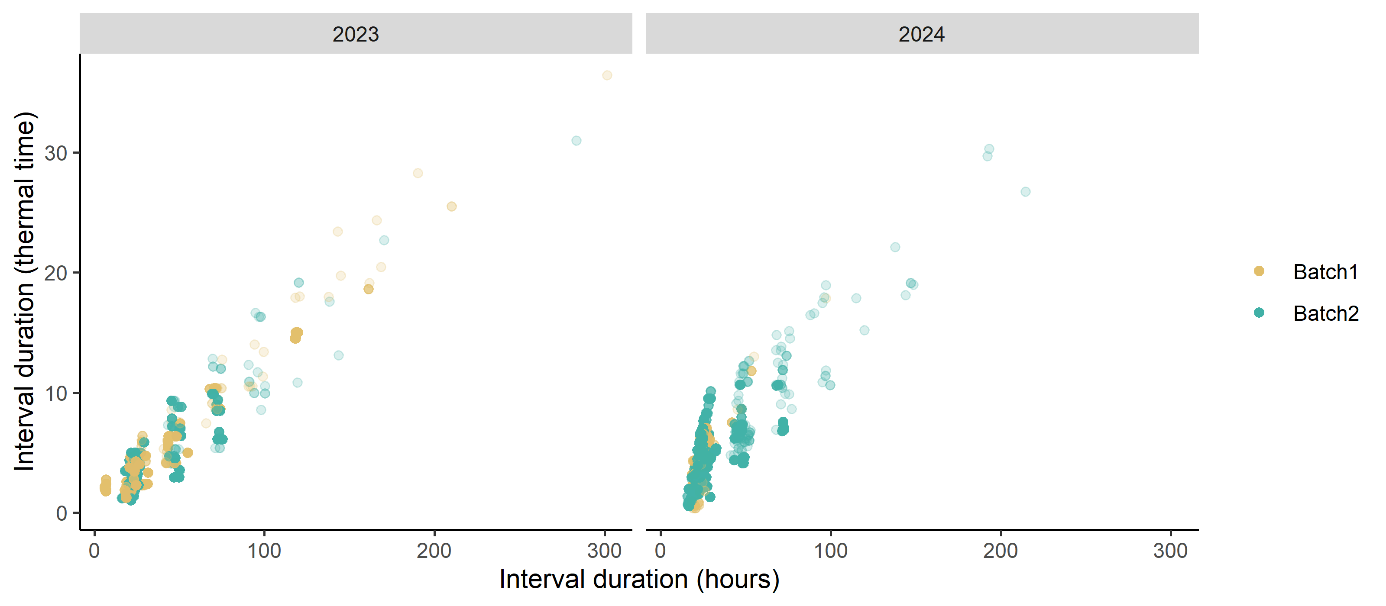

**Supplementary Figure S 6** Relationship between measurement interval duration in terms of chronological and effective thermal (temperature-corrected) time across the two years and including the two measurement batches in each experiment (Batch1 – penultimate leaves, and Batch2 – flag leaves). At constant optimal temperatures for lesion growth (estimated at 23°C; Chaloner et al., 2019), chronological time in hours is equivalent to effective thermal time. Reduced effective thermal times indicate suboptimal temperatures for lesion growth. Most measurement intervals are approximately 24 hours.

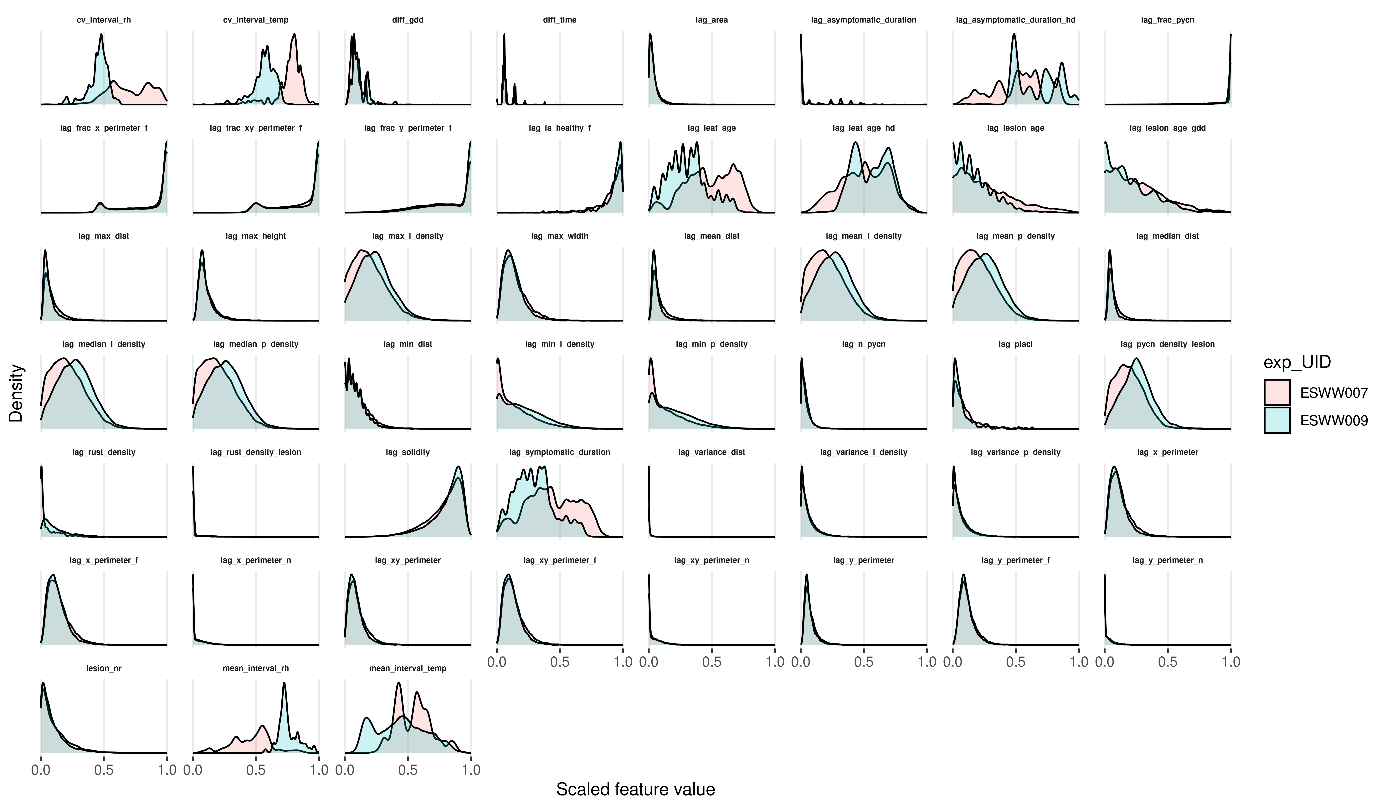

**Supplementary Figure S 7** Distribution of extracted features used for predictive modelling of lesion growth. Lesion and leaf features at the lag timepoint were used for each measurement interval. The distribution is reported separately for each experiment. Features were scaled to range from 0 to 1.

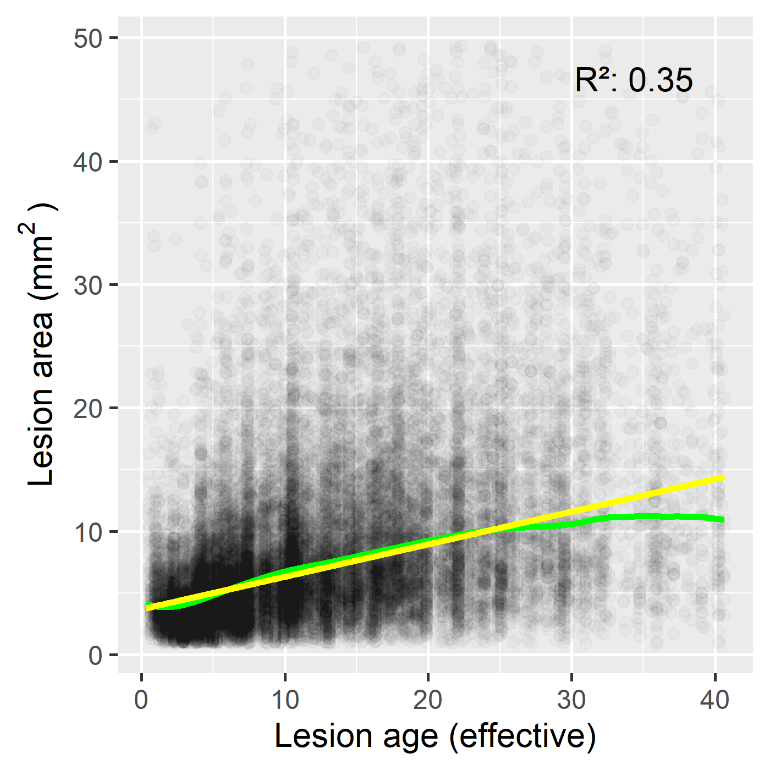

**Supplementary Figure S 8** Lesion area as a function of lesion age. The yellow line represents a linear quantile regression fit; the green curve shows a loess-smoothed trend. Axes limits were set to display all data points within the 99th percentile of lesion area and lesion age; models were fitted, and predictions are shown only within this range, due to limited data beyond this threshold. All coefficients of the linear regression model were statistically significant (p < 2.2 × 10^-16^).

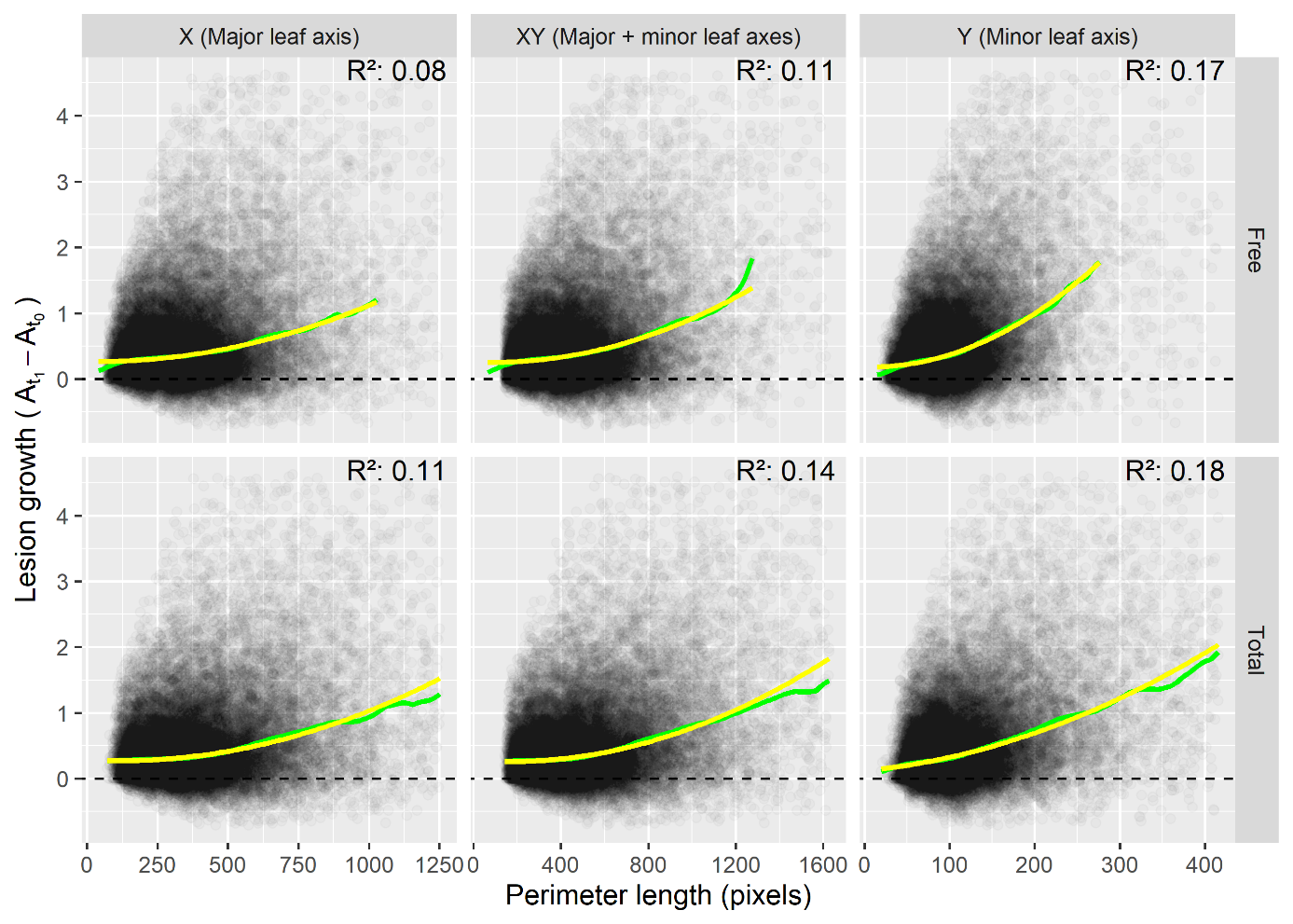

**Supplementary Figure S 9** Lesion growth vs. lesion perimeter length at t_0_ across all measurement intervals. Growth is measured as the difference in lesion area between consecutive time points ($A_{t_{1}}-A_{t_{0}}$) and is expressed per unit of thermal time. Growth is plotted against perimeter length along the major (X) and minor (Y) leaf axes, as well as their sum (XY). Top panels show unobstructed perimeters; bottom panels show total perimeters. Yellow curves represent quantile regression fits, and green lines show loess trends. X-axis limits were set to display all data points within the 99th percentile of total perimeter length values; models were fitted, and predictions are shown only within this range, due to limited data beyond this threshold (see top row). All coefficients of the second-order polynomial regression model were statistically significant (p < 2.2 × 10^-16^).

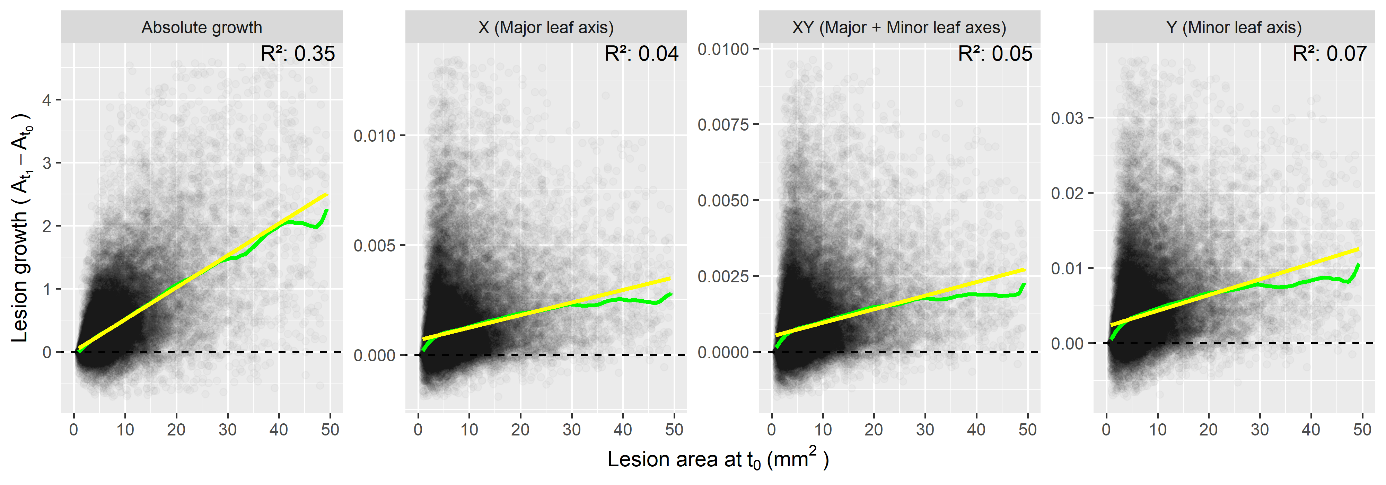
**Supplementary Figure S 10** Lesion growth as a function of lesion area at t_0_. Growth is defined as the change in lesion area between consecutive time points ($A_{t_{1}}-A_{t_{0}}$), normalized by thermal time. It is plotted as absolute growth (“None”) or per unit of lesion perimeter running along the major (X) and minor (Y) leaf axes, as well as their sum (XY). Yellow lines represent linear quantile regression fits, and green curves show loess-smoothed trends. X-axis limits were set to display all data points within the 99th percentile of lesion area values; models were fitted, and predictions are shown only within this range, due to limited data beyond this threshold. All coefficients of the linear regression model were statistically significant in all cases (p < 2.2 × 10^-16^).

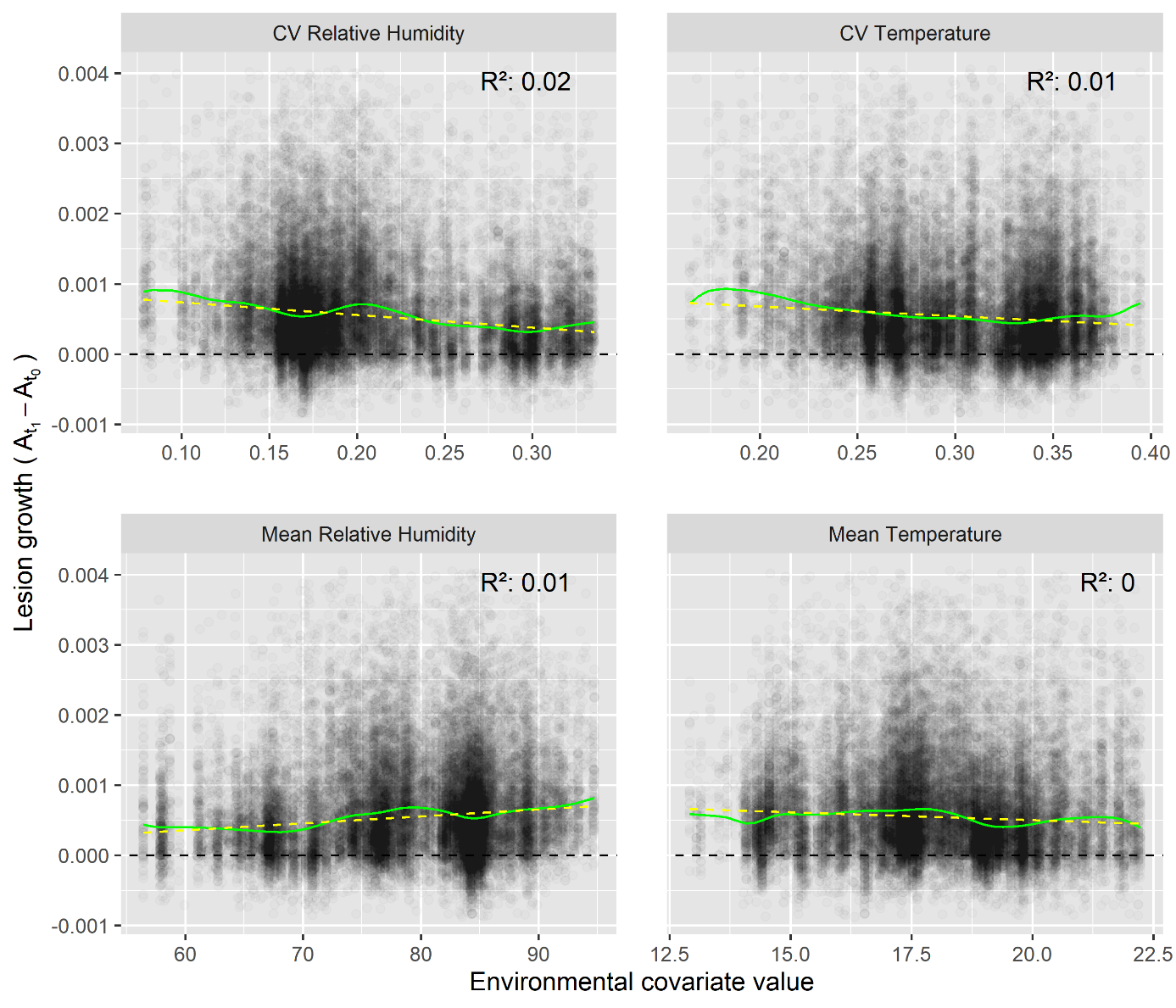

**Supplementary Figure S 11** Relationships between environmental covariates and lesion growth across all measurement intervals. Growth is measured as the difference in lesion area between consecutive time points ($A_{t_{1}}-A_{t_{0}}$) and is expressed in mm^2^ per hour. Yellow lines represent linear quantile regression fits, green lines represent loess-smoothed trends.

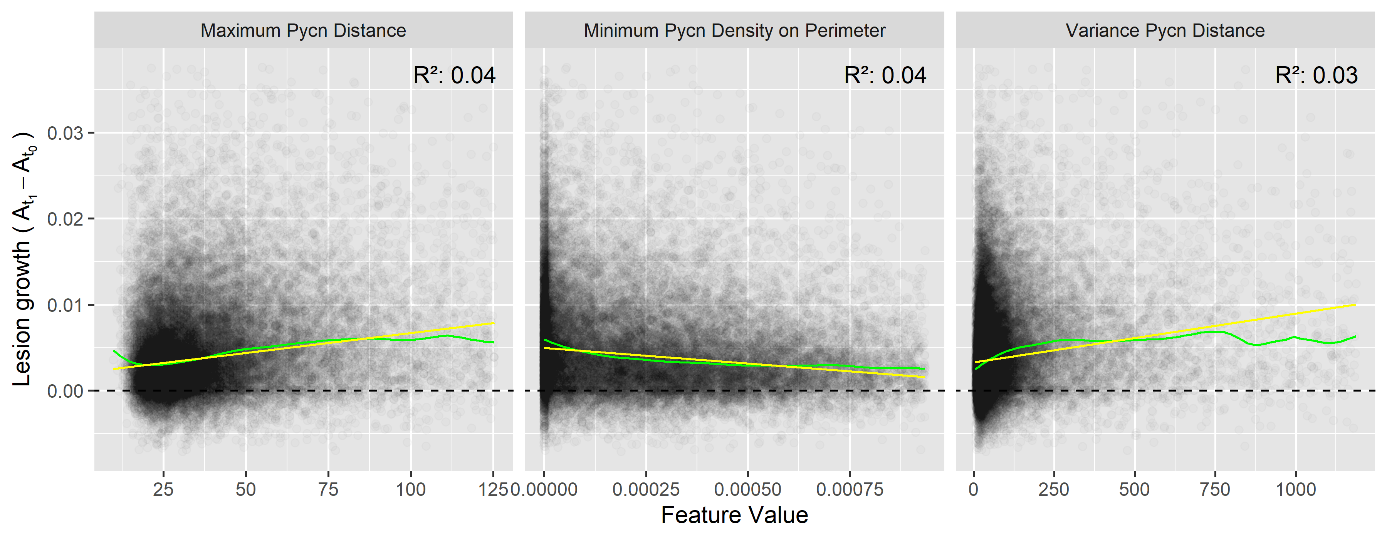

**Supplementary Figure S 12** Relationship between pycnidiation and lesion growth across all measurement intervals. Growth is measured as the difference in lesion area between consecutive time points ($A_{t_{1}}-A_{t_{0}}$) and is expressed in mm^2^ per unit of thermal time and perimeter. Yellow lines represent linear quantile regression fits, green lines represent loess-smoothed trends.

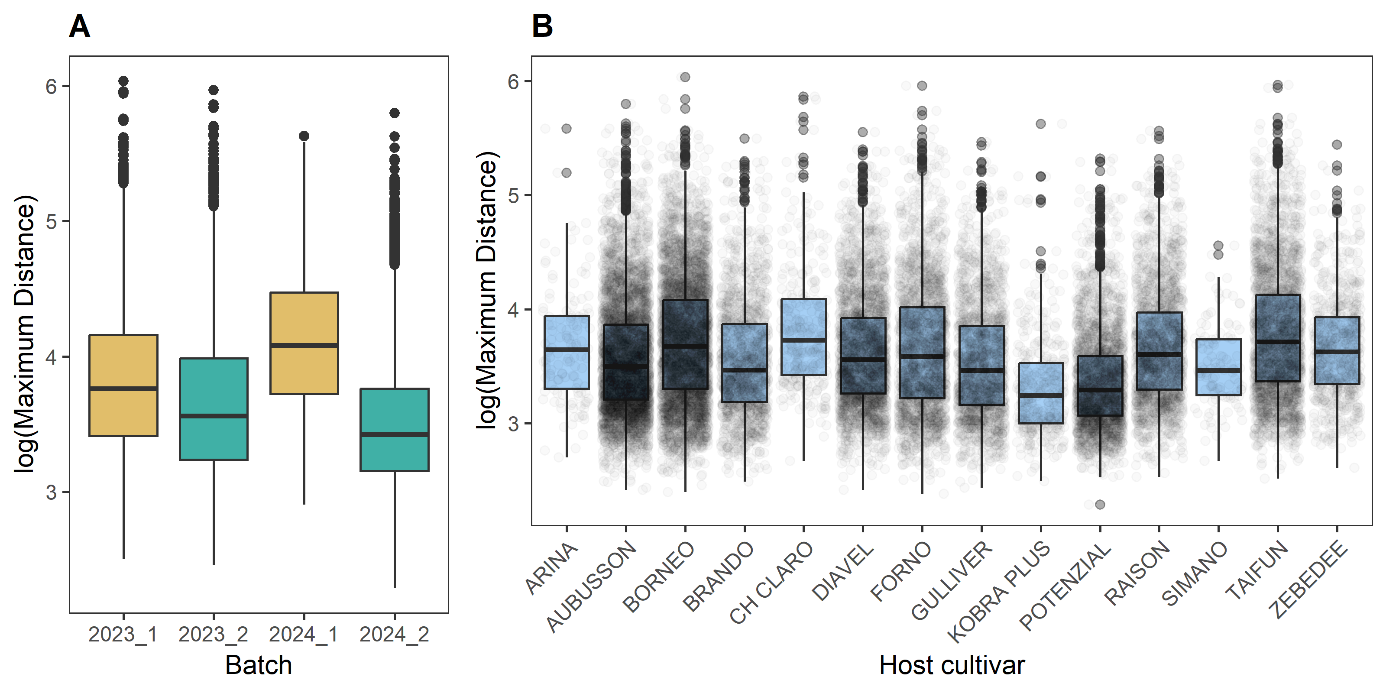

**Supplementary Figure S 13** Boxplots of the maximum distance of the lesion perimeter from pycnidia at t_0_ as related to measurement batch (**A**) and host cultivar (**B**). Distance values (zero-bound) were log-transformed to address skewness.

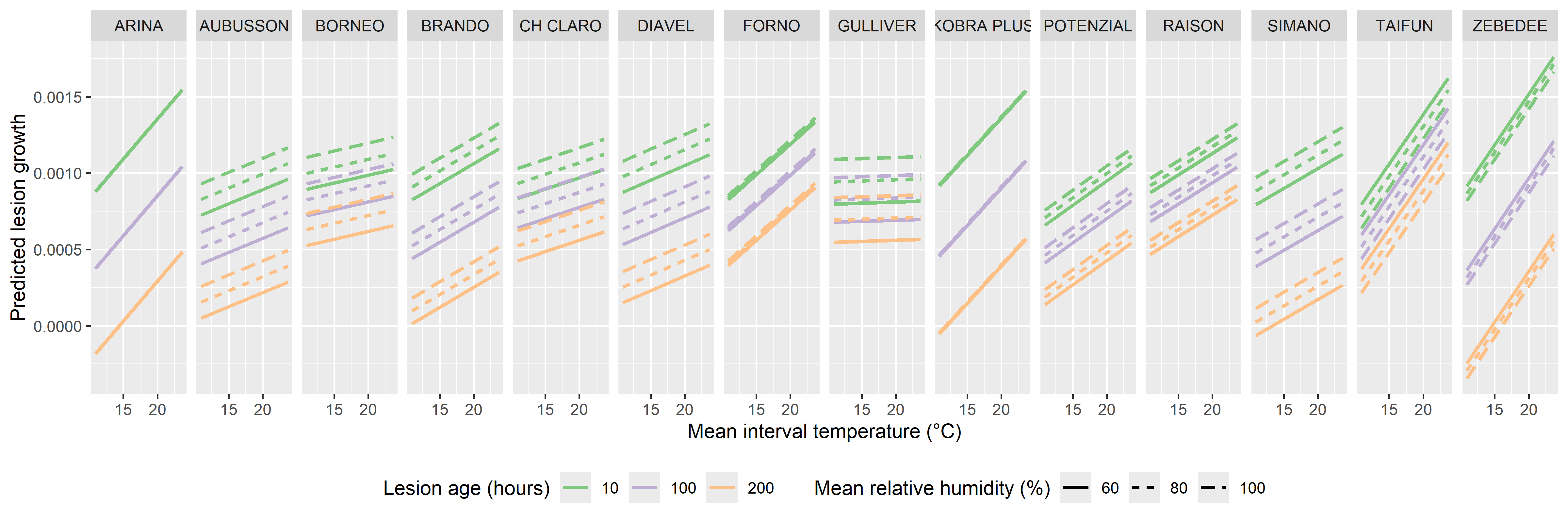

**Supplementary Figure S 14** Predicted lesion growth on 14 host genotypes over a range of mean interval temperature values, at different values of lesion age and mean interval relative humidity.

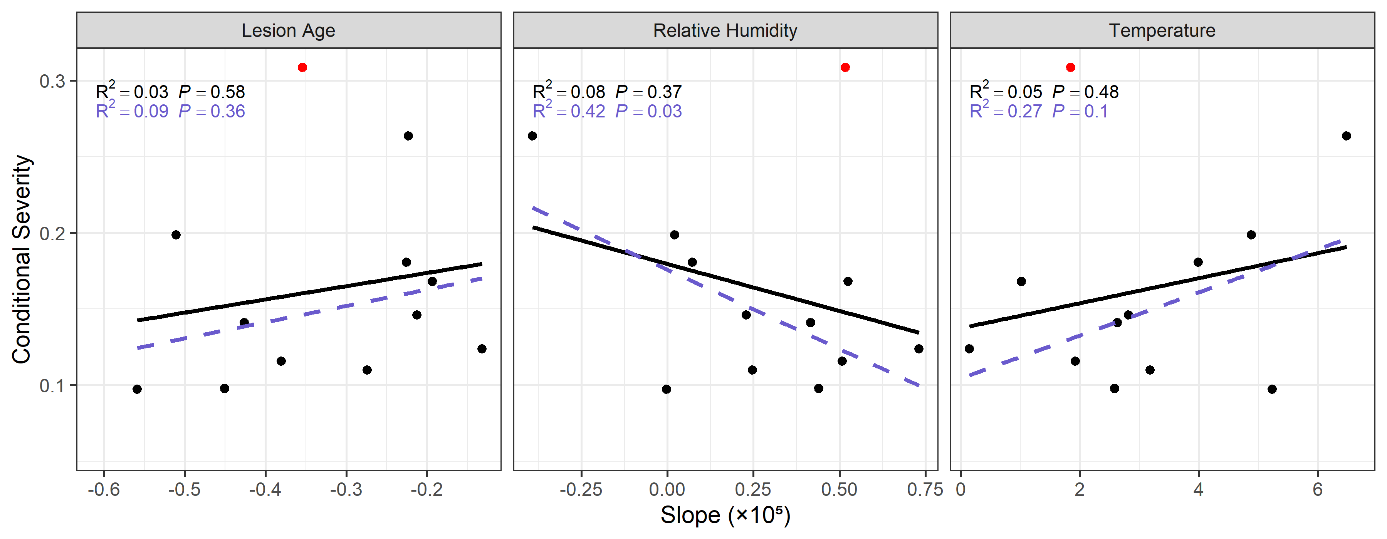

**Supplementary Figure S 15** Relationship between conditional severity and estimated linear responses of lesion growth to lesion age, mean interval relative humidity, and mean interval temperature. Genotypic BLUPs across two years are shown. Red points represent cultivar ‘AUBUSSON’. Black lines and text show linear regressions with R^2^ and *P*-values; purple lines and text show results after excluding ‘AUBUSSON’.

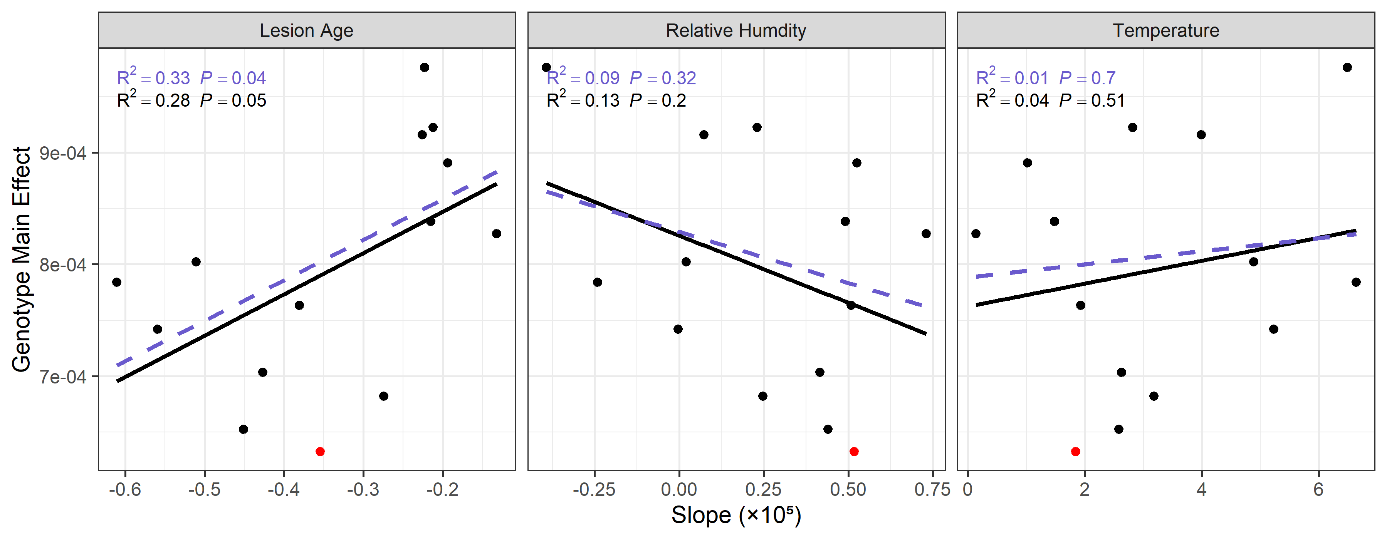

**Supplementary Figure S 16** Relationship between estimated host genotype main effects on lesion growth and linear responses of lesion growth to lesion age, mean interval relative humidity, and mean interval temperature. Genotypic BLUPs across two years are shown. Red points represent cultivar ‘AUBUSSON’. Black lines and text show linear regressions with R^2^ and *P*-values; purple lines and text show results after excluding ‘AUBUSSON’.

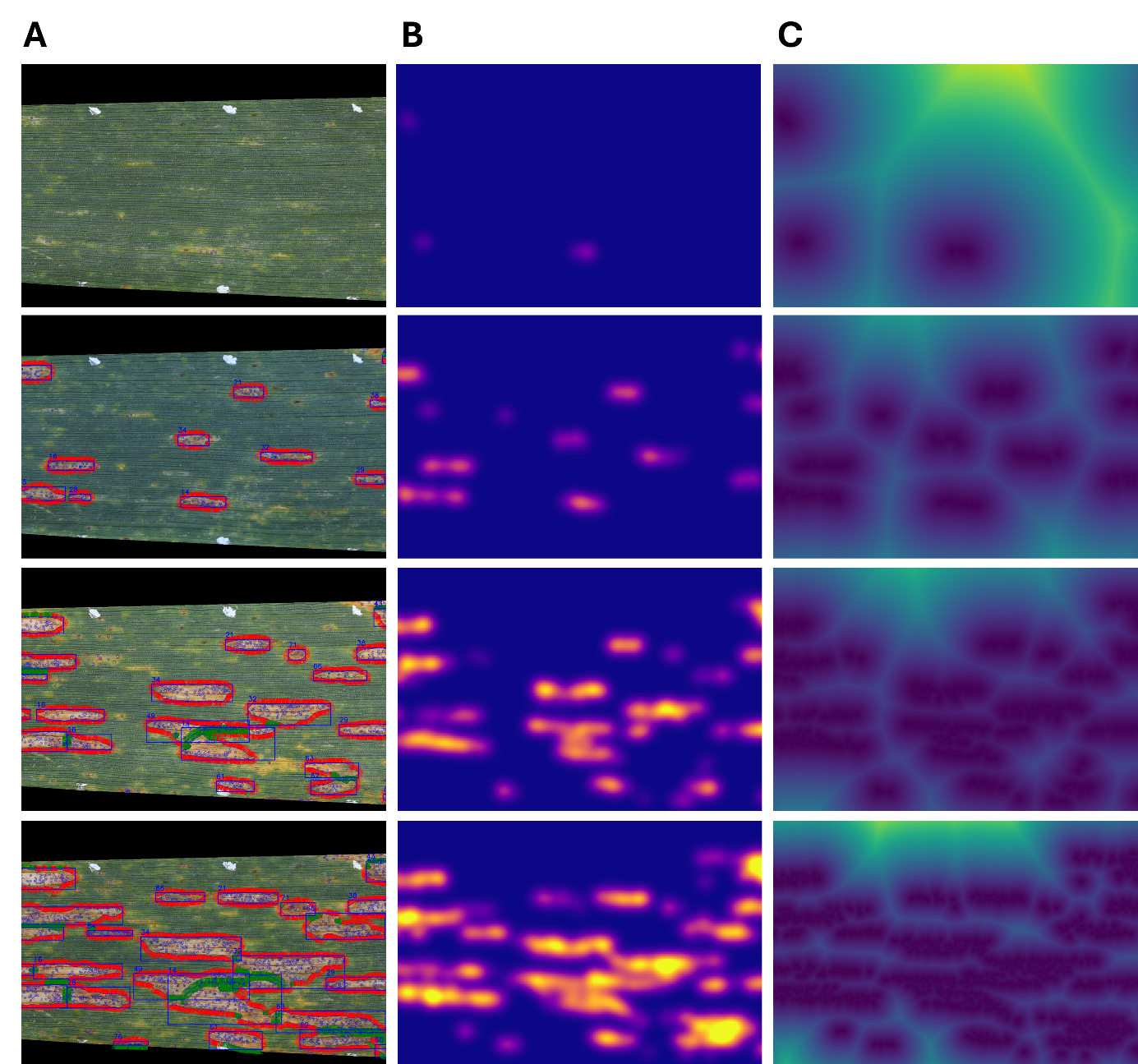

**Supplementary Figure S 17** Lesion feature extraction from base on image time series. (**A**) Cut-outs from a subset of one image series at distant time points. Lesions are marked by blue bounding boxes and attributed a number according to their sequence of appearance in the time series. Thin black lines are the b-spline approximations of the lesion perimeters, the red lines are spline normals indicating directions free of obstructions for lesion growth, whereas green spline normals indicate obstructed portions of the perimeter. Blue circles, mostly within necrotic lesions, highlight detected pycnidia and rust pustules. In these images, image axes generally align well with leaf axes, though the alignment is better for lesions located in the center of the leaf than for lesions located along the leaf edges; (**B**) Pycnidia density estimates, with brighter regions indicating higher estimated pycnidia density; (**C**) Euclidean distance transform with brighter regions indicating regions further away from pycnidia.

### Supplementary Tables

**Supplementary Table S 1** List of the wheat cultivars included in the field experiment. Cultivars were specifically selected to have similar phenology and final height but strongly contrasting canopy architectural and morphological traits, based on data from Anderegg et al. (2021). Specifically, the set comprised an equal number of cultivars with erect and planophile flag leaves and with high and low levels of flag leaf glaucousness.

| GenName | FH | GS55 | Onsen | Awns | Fl0Ang | Fl0Glc | PLACL | TraitComb | Criterion | Originator | Accession Number | Recommended for |
| --- | --- | --- | --- | --- | --- | --- | --- | --- | --- | --- | --- | --- |
| GULLIVER | 0.76 | 221 | 641 | 0 | 1 | 0 | 20.3 | LL | TraitComb | Nickerson International Research GEIE | na | GBR |
| KOBRA PLUS | 0.84 | 219 | 593 | 0 | 1 | 3 | 48.9 | LL | TraitComb | Hodowla Roslin Rolniczych Nasiona Kobierzyc Sp.z o. o. | RICP-0C0107039 | POL |
| HISTORY | 0.87 | 222 | 573 | 0 | 2 | 2 | 19.7 | LL | TraitComb | Bayerische Pflanzenzuchtgesellschaft eG & KG | RICP-01C0106532 | DEU |
| SCIROCCO | 0.92 | 218 | 587 | 0 | 5 | 2 | 32.1 | HL | TraitComb | Saatzuchtwirtschaft F. von Lochow-Petkus GmbH | RICP-01C0205033 | DEU |
| XENOS | 0.91 | 218 | 694 | 0 | 4 | 1 | 39.9 | HL | TraitComb | Fr. Strube Saatzucht | RICP-01C0106505 | AUT, LUX, DEU |
| TAIFUN | 0.85 | 216 | 656 | 0 | 5 | 3 | 37.8 | HL | TraitComb | Lochov-Petkus GMBH | K-57182;  AUS-22191 | DEU, HUN, LTU, DNK |
| RAISON | 0.75 | 222 | 579 | 0 | 1 | 8 | 29.6 | LH | TraitComb | na | na | FRA |
| BRANDO | 0.71 | 222 | 586 | 0 | 1 | 8 | 41.8 | LH | TraitComb | Cambridge PB Twyford | [K-64522](http://wheatpedigree.net/catalog/ajaxShow/K) | GBR |
| POTENZIAL | 0.82 | 221 | 699 | 0 | 2 | 9 | 22.1 | LH | TraitComb | Deutsche Saatveredelung AG, OT Leutewitz | RICP-01C0107156 | DEU, CZE, DNK |
| DIAVEL | 0.86 | 218 | 600 | 0 | 5 | 8 | ? | HH | TraitComb | Agroscope/DSP (Federal Research Station for Agronomy) | na | CHE |
| RETRO | 0.82 | 222 | 545 | 0 | 4 | 8 | 19.6 | HH | TraitComb | Nickerson Limagrain GmbH | RICP-01C0106942 | DEU |
| BORNEO | 0.74 | 221 | 523 | 0 | 6 | 7 | 42.8 | HH | TraitComb | Saatzucht J.Breun GdbR | RICP-01C0106113 | DEU |
| FORNO | 0.78 | 220 | 584 | 0 | 3 | 4 | 40.4 | na | *Lr34* | Agroscope/DSP (Federal Research Station for Agronomy) | K-62003; AUS-23641 | CHE |
| ARINA | 0.77 | 220 | 494 | 0 | 1 | 7 | 22.4 | na | Resistant | Agroscope/DSP (Federal Research Station for Agronomy) | K-57737;  K-57528; E-1014; AUS-21732 | CHE |
| MONTALBANO | na | na | na | 1 | na | na | na | na | Check | Agroscope/DSP (Federal Research Station for Agronomy) | na | CHE |
| AUBUSSON | 0.66 | 217 | 600 | 0 | 5 | 6 | 68.7 | na | Susceptible | Limagrain Verneuil Hold. | RICP-01C0107106 | FRA, ITA |
| SIMANO | 0.76 | 218 | 617 | 1 | 3 | 6 | 24.2 | na | Awned | Agroscope/DSP (Federal Research Station for Agronomy) | na | CHE |
| CH CLARO | 0.7 | 218 | 556 | 0 | 1 | 5 | 36.8 | na | Check | Agroscope/DSP (Federal Research Station for Agronomy) | na | CHE |
| ZEBEDEE | 0.7 | 222 | 624 | 0 | 3 | 9 | 28.7 | na | Other | Nickerson-Advanta Ltd. Advanta Seeds UK Ltd. | AFRC-10073 | GBR |
| CH COMBIN | 0.72 | 218 | 572 | 1 | 2 | 5 | 33.3 | na | Awned | Agroscope/DSP (Federal Research Station for Agronomy) | na | CHE |

**GenName**: Cultivar name, **FH**: Final height, **GS55**: Heading date, indicated in days after sowing; **Onsen**: Onset of senescence, indicated in growing degree days after heading; **Awns**: Presence or absence of awns on ears, with ‘0’ indicating absence and ‘1’ indicating presence of awns, **Fl0Ang**: Visual scoring of flag leaf angle, with low values indicating erect flag leaves and high values indicating drooping flag leaves, **Fl0Glc**: Flag leaf glaucousness, with low values indicating low levels of glaucousness and high values indicating high levels of glaucousness, **PLACL**: Percent leaf area covered by lesions, data from Karisto et al. (2018), **TraitComb**: Trait combination represented by the cultivar, with ‘L’ and ‘H’ representing low and high values for Fl0Ang and Fl0Glc, respectively, **Criterion**: Selection criterion applied; either the trait combination, the presence of the *Lr34* gene, or the level of resistance. Information on the originator, accession number and cultivar recommendation were taken from the Genetic Resources Information System for Wheat and Triticale (GRIS), at URL <http://wheatpedigree.net/>, accessed on 28-02-2023. Trait data was based on a field experiment carried out in the wheat growing season of 2018/2019 at the Eschikon site (Anderegg *et al.*, 2021). Assessments were made based on recommendations by Pask et al. (2012). The table was modified from Anderegg *et al.* (2023).

**Supplementary Table S 2** Description of lesion, interval, leaf, and design features evaluated as predictors of lesion growth

| **Category** | **Name** | **Description** |
| --- | --- | --- |
| lesion | area | lesion area |
|  | frac_pycn | fraction of the lesion perimeter in pycnidiation area |
|  | frac_x_perimeter_f | fraction of the free x perimeter |
|  | frac xy_perimeter_f | fraction of the free xy perimeter |
|  | frac_y_perimeter_f | fraction of the free y perimeter |
|  | lesion_age | duration between first lesion measurement and current measurement in chronological time |
|  | lesion_age_gdd | duration between first lesion measurement and current measurement in thermal time |
|  | dist | distance between perimeter and pycnidia |
|  | max_height | maximum height of lesion |
|  | max_width | maximum width of lesion |
|  | l_density | pycnidia density in lesion |
|  | p_density | pycnidia density in pycnidiation area |
|  | n_pycn | number of pycnidia in lesion |
|  | solidity | ratio of contour area to its convex hull area |
|  | x_perimeter | length of the x-perimeter |
|  | x_perimeter_f | length of the free x-perimeter |
|  | x_perimeter_n | length of the obstructed x-perimeter |
|  | xy_perimeter | length of the xy perimeter |
|  | xy_perimeter_f | length of the free xy perimeter |
|  | xy_perimeter_n | length of the obstructed xy perimeter |
|  | y_perimeter | length of the y-perimeter |
|  | y_perimeter_f | length of the free y-perimeter |
|  | y_perimeter_n | length of the obstructed y-perimeter |
|  | lesion_nr | sequential number of lesion |
| interval | diff_gdd | duration of interval in thermal time (growing degree days) |
|  | diff_time | duration of interval in chronological time |
|  | mean_interval_rh | mean interval relative humidity |
|  | cv_interval_rh | coefficient of variation of relative humidity in interval |
|  | cv_interval_temp | coefficient of variation of temperature in interval |
|  | mean_interval_temp | mean interval temperature |
| leaf | asmptomatic_duration | duration between first leaf measurement and first lesion appearing on leaf |
|  | asmptomatic_duration_hd | duration between heading and first lesion appearing on leaf |
|  | la_healthy_f | fraction of healthy leaf area |
|  | leaf_age | duration between first leaf measurement and current measurement |
|  | leaf_age_hd | duration between heading and current measurement |
|  | placl | percent leaf area covered by lesions |
|  | rust_density | rust density on leaf |
|  | symptomatic_duration | duration between first symptom detection on leaf and current measurement |
| design | exp_UID | experiment unique identifier (i.e., year) |
|  | genotype_name | host genotype name |
|  | batch_UID | leaf batch unique identifier |

**Supplementary Table S 3** Number of plots for which incidence and conditional severity were assessed at five different time points across two years. The total number of plots in the experiment was 144 and 168 in 2022 and 2023, respectively. Grey cells describe the sampled units.

| **Year** | **Time point** | **Date** | **Incidence** | | | **Conditional Severity** | | |
| --- | --- | --- | --- | --- | --- | --- | --- | --- |
|  |  |  | **L1** | **L2** | **L3** | **L1** | **L2** | **L3** |
|  |  |  | All L1&L2&L3 w/o senescence | | | All L2 w/ inc ≥ 0.33 | | |
| 2022 | t1 | 16.06.2022 | 144 | 144 | 100 | 0 | 99 | 0 |
|  |  |  | All L1 w/o senescence | | | All L1 w/ inc ≥ 0.33 | | |
| 2022 | t2 | 23.06.2022 | 140 | 0 | 0 | 144 | 0 | 0 |
|  |  |  | All L1 w/o senescence | | | All L1 w/ inc ≥ 0.33 | | |
| 2022 | t3 | 29.06.2022 | 70 | 0 | 0 | 74 | 0 | 0 |
|  |  |  | All 0F0I w/o senescence | | | all 0F0I w/ inc ≥ 0.2 | | |
| 2023 | t1 | 05.06.2023 | 54 | 54 | 50 | 31 | 49 | 46 |
|  |  |  | All L1&L2 w/o senescence | | | All L1&L2 w/o senescence | | |
| 2023 | t2 | 15.06.2023 | 168 | 166 | 0 | 70 | 118 | 0 |

**L**: Leaf layer; counting from 1 = flag leaf to 3 = flag leaf minus two, **Inc**: Incidence, assessed as a fraction of infected leaves out of a total of 30 inspected leaves; **0F0I**: No Fungicide no inoculation treatment; **w/**: with; **w/o**: without.

### Supplementary Methods

#### Lesion Feature extraction

For an exhaustive description of feature extraction, we refer to our previous publications (Anderegg *et al.*, 2022, 2024) and particularly the python code made publicly available at https://github.com/and-jonas/sympathique-wheat. Here, we provide a brief overview.

Lesion area was directly extracted as the number of pixels belonging to a connected component in the segmentation mask that was traceable to a separate infection event (cf. Supplementary Figure S17A). Area in pixels was transformed to mm^2^ by multiplying with a factor of 0.001, resulting from the sampling distance in images of approximately 0.03 mm/pixel. Features related to pycnidia density and spatial distribution were derived from (i) Euclidean distance transforms of binary masks, where pycnidia were marked as single-pixel objects, and (ii) density estimates obtained from the same masks using a Gaussian kernel with a bandwidth of 25 (Supplementary Figure S17B, C). The distance of the lesion perimeter to the nearest pycnidium and the pycnidia density along the perimeter were extracted by sampling the distance and density maps at the perimeter pixels. Summary statistics were then computed for each lesion. Within lesions, the pycnidiation area was defined as regions within 50 pixels (~1.5 mm) of a pycnidium.

Total lesion perimeter length and lesion perimeter lengths in the direction of leaf veins and perpendicular to it were approximated by summing up all differences between neighboring perimeter pixels along both image axes. Depending on the exact orientation and shape of the leaf, this approximation was occasionally not very accurate (cf. Supplementary Figure S17A). A more precise approximation would require the detection of the parallel leaf structures and adjust the measured lengths along both dimensions on a lesion-level, since not only leaf orientation, but also leaf shape affect the orientation of leaf veins in the image. This level of detail was deemed out-of-scope for the present work.

To characterize the immediate spatial context of lesions, lesion perimeters were approximated by fitting b-splines to contour pixel coordinates and sampling the segmentation masks on the corresponding spline normals (Supplementary Figure S17A). This identified portions of the perimeter with close-by obstructions to lesion growth, such as insect damage, other necrotic lesions, the leaf edge, or the edge of the region-of-interest. The corresponding absolute lengths and fractions of the lesion perimeter were extracted by summing up the length of the corresponding perimeter stretches.

#### Estimation of overall QR to STB

Detailed information on the level of QR for each cultivar included in the experiment was not available at the time of experimental design. Some genotypes were selected based on prior knowledge of their QR from (Karisto *et al.*, 2018), but that information was partial with a focus on late-season conditional disease intensity on infected leaves. Here, to obtain objective and precise estimates of QR for each genotype, we combined visual scoring of disease incidence with image-based assessments of conditional disease severity, separately for different leaf layers, following a protocol described earlier (Anderegg *et al.*, 2019). From these separate assessments, overall disease intensity is calculated by multiplying disease incidence with conditional disease severity. This strategy is very time-consuming but ideally suited to obtain reliable plot reference values for the calibration of novel phenotyping methods (Anderegg *et al.*, 2019).

Assessments were conducted on five dates across two years (2022 and 2023; Supplementary Table S3). Disease incidence was evaluated by inspecting 30 leaves per plot for visible disease symptoms. Assessments were made on all uppermost leaf layers that had not yet exhibited significant signs of physiological senescence, but never below the flag leaf minus two (i.e., for the three uppermost leaf layers at most). When multiple leaf layers were assessed, the leaves came from the same 30 culms. The resulting count data were summarized per leaf layer as a fraction of infected leaves out of the total of 30 inspected leaves. Due to the time-intensive nature of the scoring protocol, not all plots could be assessed on all dates. Consequently, data for entire leaf layers or entire treatments are missing on some dates, with focus placed on scoring parts of the experiment that provided the best discrimination among plots and genotypes. This reduced workload while preserving direct comparability across genotypes and treatments. Other missing values can be considered missing at random, unless a strong interaction between disease and phenology affected the inability to assess disease on already senescent leaves (relevant for L3, t1, 2022 and L1, t3, 2023).

Conditional severity was assessed by detaching and scanning eight infected leaves per sampled leaf layer, using conventional flatbed scanners as per (Stewart *et al.*, 2016) and segmenting the diseased area in resulting leaf images to obtain a percentage of leaf area covered by lesions as per (Zenkl *et al.*, 2025). To avoid interfering excessively with the epidemic, only plots with a disease incidence above a pre-defined threshold were sampled. The threshold was set at 0.33 in 2022 and 0.2 in 2023, reflecting the lower overall disease pressure observed in 2023 compared to 2022. Thus, in contrast to the incidence scoring, this sampling strategy most likely resulted in data missing not at random from plots with low disease incidence which is expected to correlate with conditional severity to a certain extent. Therefore, incidence and conditional severity were analyzed separately and combined to an overall estimate of QR only at the level of genotypic estimates obtained across all five assessment time points.

These overall genotypic estimates were obtained in a stage-wise approach (Piepho *et al.*, 2012; Roth *et al.*, 2021). In the first stage, a spatial correction model was fitted to plot-based disease incidence and conditional severity values separately, for each time point. The model included a spatially structured random effect modeled via a two-dimensional P-spline over plot design coordinates, as well as random effects for row and column. Given the repeated measurement characteristic of sampling multiple leaf layers per plot, a plot unique identifier was included in the random term. Fixed effects accounted for treatment, leaf layer, scorer, and block (replicate). Genotype was treated as a fixed effect to obtain best linear unbiased estimates (BLUEs), or as a random effect to obtain an estimate of within-timepoint heritability. Both incidence and conditional severity data were logit-transformed to obtain a normal distribution. Accordingly, a gaussian error distribution was used for both traits. For conditional severity, a vector of weights (*w*) was included in the fitting process based on error variance estimations as *w* = 1/(*var*(*e*)), with *var*(*e*) denoting the variance of PLACL in each plot, estimated for the sample of 8 leaves (Roth *et al.*, 2021). Thus, the full model was:

$$\begin{aligned} Y_{ijklmn}=f\left( r_{i}, c_{j} \right)+R_{i}+C_{j}+P_{k}+G_{l}+T_{m}+S_{n}+L_{o}+\varepsilon_{ijklmno}\#\left( 4 \right) \end{aligned}$$

where $f\left( r_{i}, c_{j} \right)$ is a smoothed bivariate surface defined by row r (i = 1, …,14) and range c (j = 1, …, 15), $R_{i}$ and $C_{j}$ are random factors of the rows and ranges, respectively, $P_{k}$ is the random effect of the k^th^ plot, $G_{l}$ is the fixed effect of the l^th^ genotype, $T_{m}$ is the fixed effect of the m^th^ treatment (m = 1, …, 4), $S_{n}$ is the fixed effect of the n^th^ scorer (n = 1, …, 4), $L_{o}$ is the fixed effect of the o^th^ leaf layer (o = 1, 2, 3), and ε is the residual error. The model was fitted using the R-Package ‘SpATS’ (Rodriguez-Alvarez *et al.*, 2019). The model was reduced when factors were not relevant (e.g., when only one leaf layer was scored, the fixed effect of the leaf layer and the random plot effect were omitted). This first stage yielded a genotypic BLUE for incidence and conditional severity for each cultivar for each time point. These were processed in the second stage to obtain an overall estimate of genotype effects and heritability across all time points, still separately for incidence and conditional severity, using the linear mixed model:

$$\begin{aligned} Y_{lt}=\mu+T_{t}+G_{l}+ {GT}_{tl}+\varepsilon_{lt}\#\left( 5 \right) \end{aligned}$$

where $Y_{lt}$ is the spatially adjusted genotype mean (BLUE), $\mu$ is the overall mean, $T_{t}$ is the fixed effect of the t^th^ time point (t = 1, …, 5), $G_{l}$ is the random effect of the l^th^ genotype, ${GT}_{tl}$ is the random genotype-by-timepoint interaction, and ε is the residual error. A vector of weights (*w*) was included as *w* = 1/se^2^, with se denoting the standard errors of the BLUEs from the first stage. This model allowed the estimation of variance components to calculate heritability across time points and provided an overall estimate of the genotype effects. The model was fitted in R using ASReml-R (Butler *et al.*, 2018).

### References

**Anderegg J, Aasen H, Perich G, Roth L, Walter A, Hund A**. **2021**. Temporal trends in canopy temperature and greenness are potential indicators of late-season drought avoidance and functional stay-green in wheat. *Field Crops Research* **274**: 108311.

**Anderegg J, Hund A, Karisto P, Mikaberidze A**. **2019**. In-Field Detection and Quantification of Septoria Tritici Blotch in Diverse Wheat Germplasm Using Spectral–Temporal Features. *Frontiers in Plant Science* **10**.

**Anderegg J, Kirchgessner N, Kronenberg L, McDonald BA**. **2022**. Automated Quantitative Measurement of Yellow Halos Suggests Activity of Necrotrophic Effectors in Septoria tritici Blotch. *Phytopathology®* **112**: 2560–2573.

**Anderegg J, Zenkl R, Kirchgessner N, Hund A, Walter A, McDonald BA**. **2024**. SYMPATHIQUE: image-based tracking of symptoms and monitoring of pathogenesis to decompose quantitative disease resistance in the field. *Plant Methods* **20**: 170.

**Anderegg J, Zenkl R, Walter A, Hund A, McDonald BA**. **2023**. Combining High-Resolution Imaging, Deep Learning, and Dynamic Modeling to Separate Disease and Senescence in Wheat Canopies. *Plant Phenomics* **5**: 0053.

**Butler DG, Cullis BR, Gilmour AR, Gogel BJ, Thompson R**. **2018**. ASReml estimates variance components under a general linear mixed model by residual maximum likelihood (REML). R Package version 4.2.0.355. : 188.

**Chaloner TM, Fones HN, Varma V, Bebber DP, Gurr SJ**. **2019**. A new mechanistic model of weather-dependent Septoria tritici blotch disease risk. *Philosophical Transactions of the Royal Society B: Biological Sciences* **374**: 20180266.

**Karisto P, Hund A, Yu K, Anderegg J, Walter A, Mascher F, McDonald BA, Mikaberidze A**. **2018**. Ranking Quantitative Resistance to Septoria tritici Blotch in Elite Wheat Cultivars Using Automated Image Analysis. *Phytopathology* **108**: 568–581.

**Pask A, Pietragalla J, Mullan D, Reynolds MP**. **2012**. Physiological breeding II : a field guide to wheat phenotyping. : iv, 132 pages.

**Piepho H-P, Möhring J, Schulz‐Streeck T, Ogutu JO**. **2012**. A stage‐wise approach for the analysis of multi‐environment trials. *Biometrical Journal* **54**: 844–860.

**Rodriguez-Alvarez MX, Lee D-J, Kneib T, Durban M, Eilers P**. **2019**. *SAP: Multidimensional Generalized P-Splines Regression Models Estimation*.

**Roth L, Rodríguez-Álvarez MX, van Eeuwijk F, Piepho H-P, Hund A**. **2021**. Phenomics data processing: A plot-level model for repeated measurements to extract the timing of key stages and quantities at defined time points. *Field Crops Research* **274**: 108314.

**Stewart EL, Hagerty CH, Mikaberidze A, Mundt CC, Zhong Z, McDonald BA**. **2016**. An Improved Method for Measuring Quantitative Resistance to the Wheat Pathogen Zymoseptoria tritici Using High-Throughput Automated Image Analysis. *Phytopathology* **106**: 782–788.

**Zenkl R, McDonald BA, Walter A, Anderegg J**. **2025**. Towards high throughput in-field detection and quantification of wheat foliar diseases using deep learning. *Computers and Electronics in Agriculture* **232**: 109854.
